# Ancient DNA resolves the taxonomic status and functional characteristics of the Pleistocene lowland yak (*Bos baikalensis*)

**DOI:** 10.64898/2026.11.23.768568

**Authors:** Alexandre Gilardet, Jonas Oppenheimer, Maxim B. Kozlikin, Nathan Martin, Georgios Xenikoudakis, Michael V. Shunkov, Katerina Douka, Sergey K. Vasiliev, Peter D. Heintzman, Love Dalén

## Abstract

Species delineation and taxonomy are notoriously complicated, especially for fossils where available information is often limited to morphology. However, palaeogenomics can add a genetic dimension to fossils, which allows for both robust resolution of taxonomy and trait functional characterization. In this study, we present the two first ancient genomes from voucher specimens morphologically attributed to the extinct yak *Bos mutus baikalensis*. We demonstrate that this taxon falls within the genetic variation of yak X, a cryptic bovine species previously identified using genetics solely from non-diagnostic bone fragments. We collate morphological, palaeontological, geographical, and genomic data available from both *Bos mutus baikalensis* and yak X in order to fully describe this species, which we propose be named as the lowland yak (*Bos baikalensis*). We further characterize genomic regions that may have been under natural selection or underlie functional differences in the lowland yak compared to the extant wild mountain and domestic yaks (*B. mutus* and *B. grunniens*, respectively) and other bovines. We find signals of functional changes in genes related to hypoxia, appetite control, morphology, fat metabolism, temperature, olfaction, immunity, and reproduction.

## Introduction

The species concept has many, sometimes incompatible, definitions. The conflict between definitions stems in part from the complexity and diversity of the data used to observe and describe biological entities. With Linnaeus, earlier classifications relied on phenetics, or the phenotypic quantitative characteristics of organisms (Sokal and Crovello 1970). Others have emphasized ecological niches (Van Valen 1976), or phylogenetic approaches relying on monophyly or shared genetic variation (D. E. Rosen 1979; Donoghue 1985; Baum and Shaw 1995). A palaeontological perspective additionally demonstrated the temporal and evolutionary dimension of a species, or lineage (Simpson 1961). De Queiroz (de Queiroz 2005) suggested the need for an integrated approach that relies on multiple such definitions, depicting speciation as a gradual process eventually satisfying most definitions of the species concept, in no defined order.

Fossils are a valuable resource for understanding the evolution of taxonomic groups, as well as to organize biological diversity into ancestors and descendants. Palaeontology has historically relied on morphological characteristics, morphometrics, geography, and ecology (*i.e.* isotopes) to delineate taxa. Methodological advances in the recovery of organic material, including ancient DNA (aDNA), from palaeontological materials (Korlević et al. 2015; Burrell et al. 2015; Damgaard et al. 2015; Leonardi et al. 2017) now allow access to another layer of information from fossils (Eisenmann and Vasiliev 2011). Palaeogenomics has helped characterize several new lineages previously unknown from the Quaternary palaeontological record, such as the Denisovan hominin (Reich et al. 2010) or the Krestovka mammoth (van der Valk et al. 2021), and has proposed new taxonomic classifications, such as assigning the North American stilt-legged horse to a new genus *Haringtonhippus* (Heintzman et al. 2017).

Recent genomic work has identified a new bovine species in Pleistocene mid-latitude Asia, termed yak X, that is deeply divergent from the extant wild yak (*Bos mutus*), which we refer to as ‘mountain’ yak hereafter, and domestic yak (*B. grunniens*) (Gilardet et al. 2025; Oppenheimer, Gilardet, et al. 2026). Yak X was identified from non-diagnostic bone fragments or partially diagnostic bones, making it challenging to designate an appropriate scientific name for this species. However, the Baikal yak (*Bos mutus baikalensis*), currently recognised from palaeontological descriptions as a larger extinct subspecies of the extant yak in the region (Vasiliev 2021), was proposed as a potential candidate (Oppenheimer, Gilardet, et al. 2026). To resolve whether yak X corresponds to the Baikal yak, we generated genomic data from specimens unequivocally identified as Baikal yak on the basis of morphology, and compared them to the yak X genomes, showing that they are the same taxon that we call the lowland yak (*Bos baikalensis* Verestchagin 1954 (Verestchagin 1954)). The mid-latitude Asian sites where lowland yak has been detected differ in a number of ways from the Qinghai–Tibet Plateau (QTP) where the ancestors of extant yaks evolved (K. Zhang et al. 2020), including being at lower elevation and having better food availability (Vasiliev 2021), suggesting potential functional changes between them. Through an integrative approach combining palaeontological, morphological, geographical, and genomic information, we refine the taxonomy of the lowland yak, and functionally characterize it.

## Results

### The Baikal yak and yak X are the same genetic lineage

We generated genomic data from two voucher specimens morphologically identified as Baikal yak, which placed these individuals within yak X phylogenetic diversity. The two Baikal yak voucher specimens were excavated from two sites in southern Siberia, consisting of a metacarpal (AG279) from Denisova Cave and a humerus (AG281) from Fanatikov Cave, and were taxonomically assigned based on morphological characteristics and measurements (Vasiliev 2021) (Figs. 1A, S1; Table S1; Supplementary Information).

**Figure 1:**
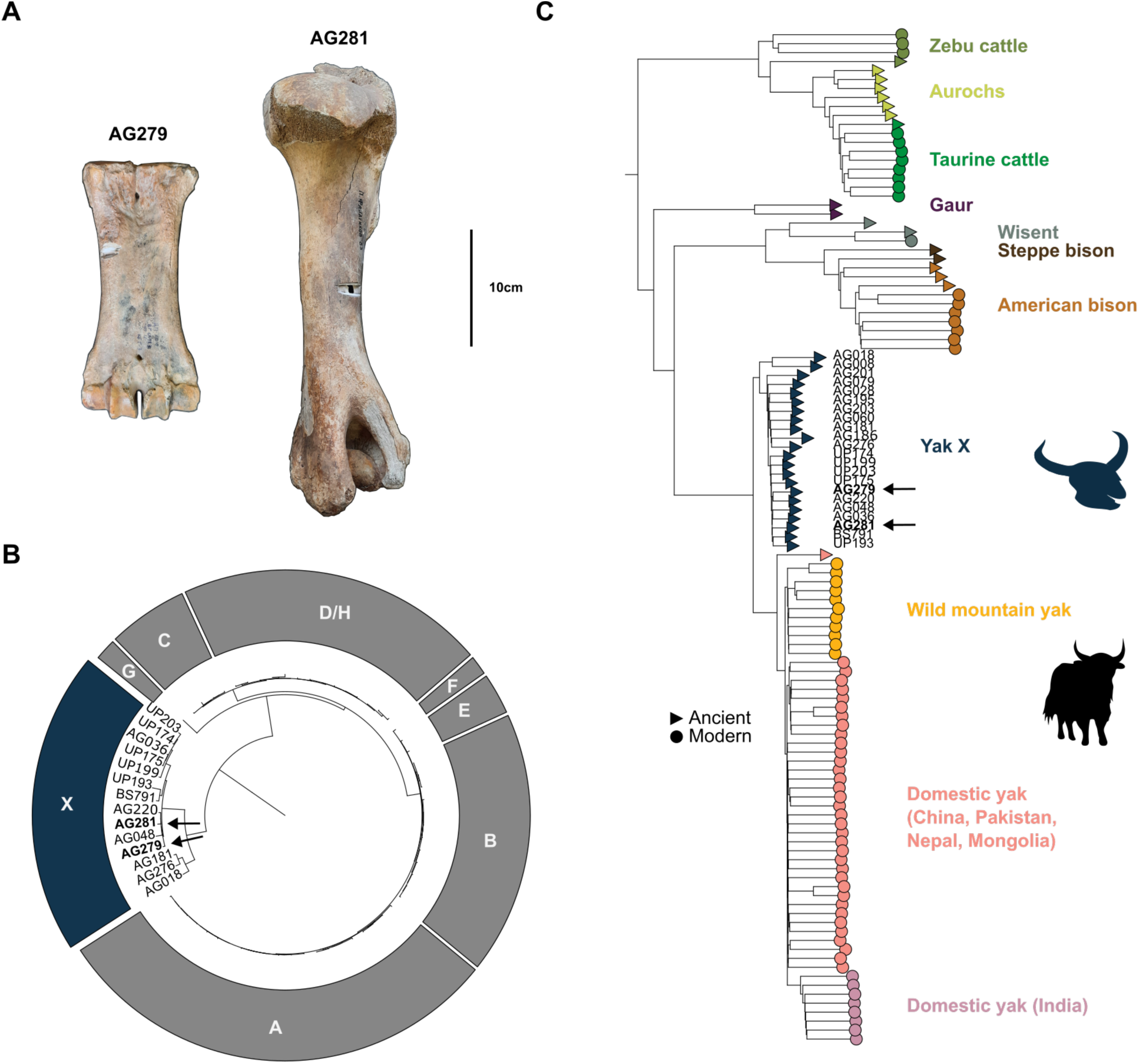
Morphology and phylogenetic placements of the voucher Baikal yak specimens. A) Metacarpal (AG279 - collection ID: DCE-10) and humerus (AG281) morphologically identified as Baikal yak and recovered from southern Siberia. Scale is shown. Additional pictures are available in Fig. S1. B) Maximum-likelihood mitochondrial and C) neighbour-joining autosomal phylogenies of the two Baikal yak voucher specimens, denoted in bold and with an arrow. Tip labels are B) modern yak and yak X haplogroups or C) Bos and Bison individuals. Support and all individual IDs are shown in Figs. S2 and S3. Trees are rooted with B) bison or C) water buffalo (not shown).

We sequenced between ∼238 and 262 million sequencing reads to generate 41-fold and 44-fold depth of coverage of the mitogenomes, as well as 0.2-fold and 0.8-fold depth of coverage of the autosomal genomes, for AG279 and AG281, respectively. In both the mitochondrial and autosomal phylogenies, the two Baikal yak voucher specimens cluster within yak X genetic diversity (Figs. 1B, 1C), with 100% bootstrap support in both phylogenies (Figs. S2, S3). More specifically, both Baikal yak specimens cluster within previously described Late Pleistocene yak X diversity, in mitochondrial subclade 2 and autosomal subclade 4b (Oppenheimer, Gilardet, et al. 2026). A principal component analysis (PCA) containing yak diversity additionally supports the placement of the voucher Baikal yak specimens within yak X diversity (Fig. S4). We additionally calculated D-statistics to show that the two voucher Baikal yak specimens do not have affinity to other bovine species — wild mountain yak, bison (*Bison* sp.), and aurochs (*Bos primigenius*) — relative to yak X (Fig. S5A-C), and to demonstrate excess allele sharing between Baikal yak and yak X compared to wild mountain yak (Fig. S5D). Overall, we confidently demonstrate that Baikal yak and yak X are the same species, *Bos baikalensis* Verestchagin 1954. Given its wider distribution beyond the areas surrounding Lake Baikal, we suggest to commonly refer to this extinct species as the ‘lowland’ yak to contrast with its closest known relative, the higher elevation-dwelling extant wild ‘mountain’ yak.

Genomic sexing of both voucher specimens is in agreement with previous morphological assignments of the metacarpal (AG279) as male and the humerus (AG281) as female. These findings lend further support to palaeontological studies suggesting that sexual dimorphism was more pronounced in the lowland yak compared to steppe bison (*Bison priscus*) (Vasiliev 2021). Together with the observed sexual dimorphism in wild mountain yaks where a female can be one-third of the mass of a male (Leslie and Schaller 2009), we hypothesize that increased sexual dimorphism might have evolved in the common ancestor of both wild mountain and lowland yaks. This could be due to increased sexual competition in a polygynous mating system, consistent with female mate choice selecting larger size males in wild mountain yaks (Buzzard et al. 2014).

### Geographical range of the Pleistocene lowland yak

We collated all locations of both Baikal yak (Flerow 1980; Vasiliev 2021) and yak X (Oppenheimer, Gilardet, et al. 2026) fossil finds to refine the overall geographical range of the lowland yak (Fig. 2; Table S2). While the extant wild mountain yak is restricted to the high altitudes of the QTP (Schaller 1998; Leslie and Schaller 2009), the lowland yak ranged from the Siberian Altai and Khakassia regions to Transbaikalia across the Altai, Sayan and Khangai Mountains, thus stretching from Russian Siberia into Mongolia. We also highlight two outliers of the lowland yak’s identified main range in Nizhnyaya Tavda, East of the Ural Mountains, with a non-finite radiocarbon date (Gilardet et al. 2025) and in Central Yakutia, by the Vilyuy River, with no available chronological information (Flerow 1980; Vasiliev 2021). We note that the latter outlier echoes a reported putative yak further north in Chukotka (Flerow 1980). The QTP and the range inhabited by the Pleistocene lowland yak are separated by the east-west Tian Shan Mountain and neighbouring low-altitude semi-continuous deserts and arid areas. The broad Gobi Desert region (Heiner et al. 2019) connected to the Tarim Basin and its central Taklamakan Desert lie on the east side of the Tian Shan, while several neighbouring deserts lie on the west side, including the Muyunkum, Karakum and Kyzylkum Deserts of Central Asia (W. Wang et al. 2020). These topographic and environmental elements, partially in place since at least the Early Pleistocene (X. Wang et al. 2016; J. Sun et al. 2017; Lu et al. 2019), were already suggested as possible geographical and ecological barriers that resulted in the observed divergence between mountain and lowland yaks (Oppenheimer, Gilardet, et al. 2026), and are concordant with patterns observed in other species (Mahmut et al. 2002; Lebedev et al. 2021; Suzuki et al. 2023).

**Figure 2:**
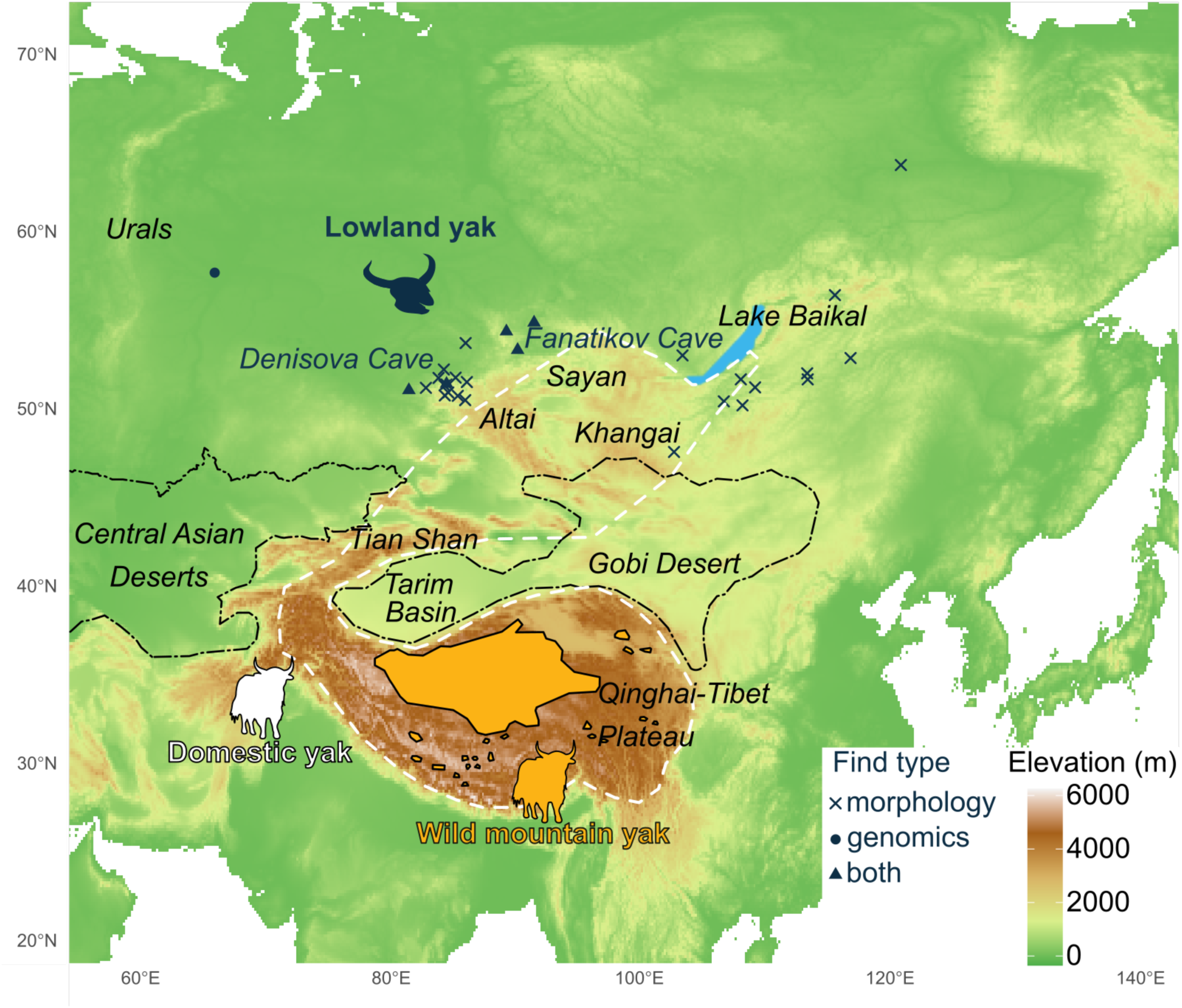
Geographical ranges of the Pleistocene lowland, wild mountain and domesticated yaks. Sites are indicated by: crosses where the Baikal yak was morphologically identified; filled circles where yak X was solely genetically identified; and triangles where both Baikal yak and yak X were identified, including Denisova Cave and Fanatikov Cave where we report morphological and genetic identification of the same bones. The present-day distribution of wild mountain yak and domestic yak are shown in orange and with a white dashed line, respectively (Schaller 1998; Leslie and Schaller 2009; K. Zhang et al. 2020). Desert regions are delineated with black dash-dotted lines. The name and extended information for each plotted site are available in Table S2.

Domestic yaks (*Bos grunniens*), derived from the wild mountain yak (Qiu et al. 2015), were introduced beyond the QTP into neighbouring regions of China, Bhutan, India, Nepal, Kyrgyzstan, Kazakhstan, Tajikistan, Afghanistan and Pakistan, as well as Mongolia and Siberia (C. Li et al. 2003; Xuebin et al. 2005; K. Zhang et al. 2020) (Fig. 2), thus overlapping geographically, but perhaps not temporally, with the extinct Pleistocene lowland yak, as the earliest yak domestication estimate is dated to 7,300 years before present (Qiu et al. 2015). However, we note that rock art from the Mongolian Altai has been suggested to represent the co-occurrence of wild and domestic yaks (Jacobson-Tepfer 2025), and historical reports of wild yaks in Tuva, by the Altai-Sayan (Flerow 1980; Vasiliev 2021) and in Dauriya, Transbaikalia (Flerow 1980) could suggest a survival of the lowland yak much later into the Holocene (Vasiliev 2021). Since a previous microsatellite-based study suggested a significant genetic differentiation of Russian and Mongolian domestic yaks compared to a QTP domestic yak population (Xuebin et al. 2005), we specifically tested for allele sharing between the lowland yak and two Mongolian yaks as compared to other domestic yaks. We do not detect affinities between Mongolian domestic yaks and lowland yaks (Fig. S6), although we caution that this analysis could benefit from additional Russian and Mongolian yak individuals.

### Functional changes in the Pleistocene lowland yak

Pleistocene environments in the Altai-Sayan-Khangai Mountains, Transbaikalia and neighbouring steppe differed from those of the QTP. The latter was characterized by higher altitude, lower oxygen levels, and scarcer food availability (Vasiliev 2021). These distinct environmental conditions, along with the deep genomic divergence between lowland and mountain yaks, and the apparent lack of gene flow between them after ca. 250,000 years ago (Oppenheimer, Gilardet, et al. 2026) suggest the possibility that specific adaptations would have evolved in the lowland yak. To investigate this, we used a version of the Population Branch Statistic (PBS) based on *f3*-statistics to identify highly divergent regions in twelve lowland yak genomes (including AG279 and AG281), relative to mountain yak and American bison, possibly indicative of selective sweeps. Depending on the specific yak and bison used as outgroups in the PBS test (Figs. 3A, S7; Table S3), we find eleven regions that were consistently and significantly differentiated in the lowland yak compared to these other groups, suggesting the presence of a small number of hard selective sweeps. The region on chromosome 1 overlaps *RCAN1*, a gene involved in oxidative stress and shown to interact with HIF-1α (a Hypoxia Inducible Factor) (Yun et al. 2022). *RCAN1* was highlighted in the high-altitude adaptation of a primate, the gelada (*Theropithecus gelada*) (Chiou et al. 2022), thus potentially pointing at an altitude-related functional change in the lowland yak. Among other regions, we point out two that occur in olfactory gene clusters, and a potential immunity-related signal with *WC1* (Baquero and Plattner 2017), several guanylate-binding proteins (*GBPs*) previously linked to immunity and growth traits in cattle (Hao et al. 2020; M. Chen et al. 2026), as well as *MRP4*, linked to both reproduction and inflammation (Lacroix-Pépin et al. 2011; M. T. Lawrence et al. 2026).

**Figure 3:**
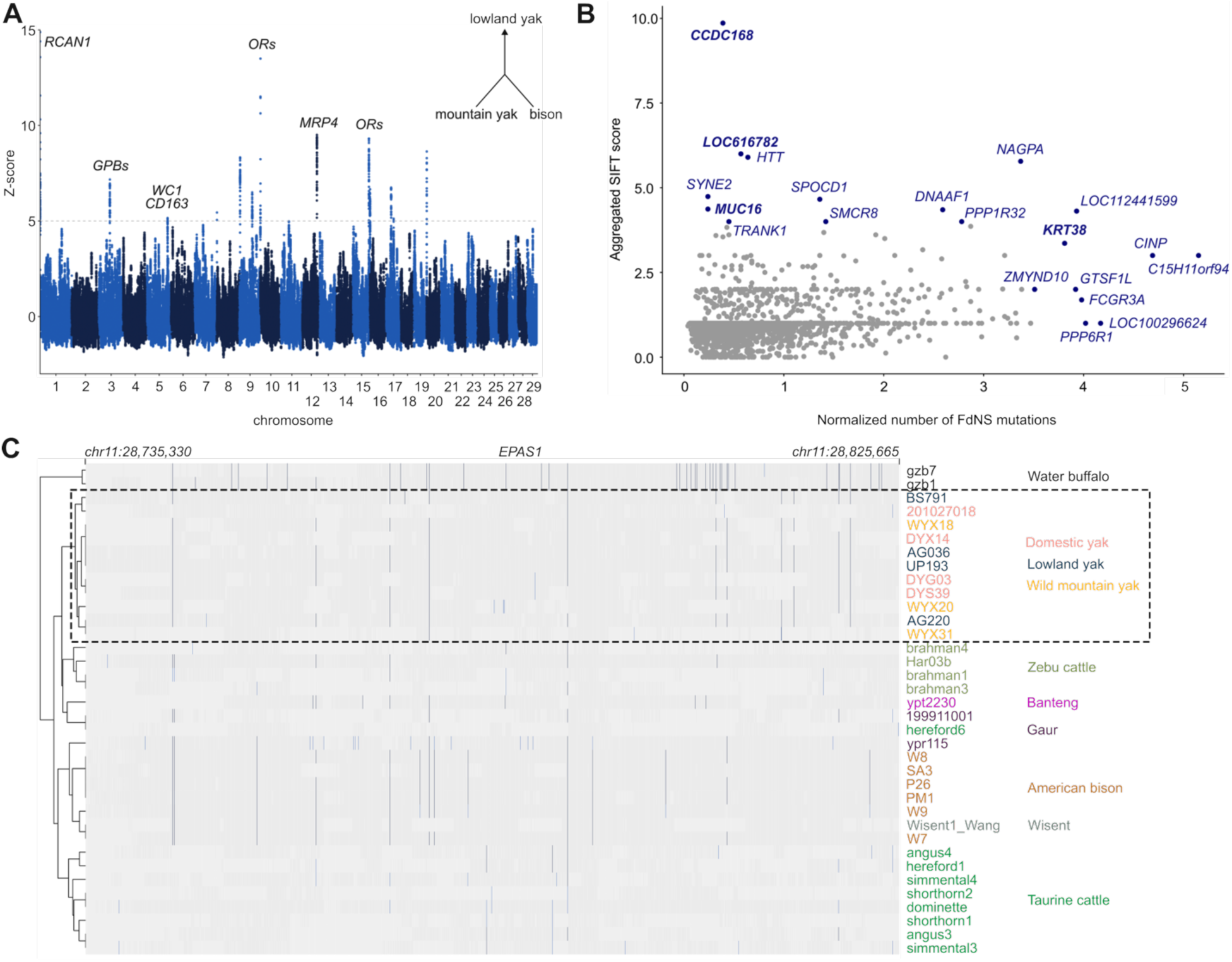
Functional changes in the lowland yak. A) f3-statistics-based Population Branch Statistic (PBS) outlier test, identifying highly differentiated regions in the lowland yak relative to wild mountain yak and bison. A schematic of the statistics is depicted in the top right corner. RCAN1 (Regulator of Calcineurin 1), GPBs (Guanylate-Binding Proteins), ORs (Olfactory Receptors), MRP4 (Multidrug-Resistance-Associated Protein 4). Details on windows and overlapping genes are in Table S3. Alternative versions of the test are shown in Fig. S7. B) Genes containing possible FdNS mutations plotted relative to their aggregated SIFT score and number of mutations normalized by gene length (Table S4). Genes with a SIFT score or normalized number of mutations above 4 and 3.5, respectively, are shown in blue, with the ones we discuss in the main text highlighted in bold. C) Heatmap of the shared derived state of the EPAS1 gene across all yaks in the context of Bos and Bison. Only exons are shown. Homozygous alternative sites are shown in darker blue, heterozygous sites are shown in lighter blue, homozygous reference sites are shown in darker grey and missing sites are shown in lighter grey. A phylogenetic tree is shown in Fig. S8.

We identified genes containing possible fixed-derived non-synonymous (FdNS) mutations by comparing four published lowland yak genomes (>3-fold depth of coverage) relative to wild mountain yak and bison (Fig. 3B). We detected 1,564 genes (1,802 when including uncharacterized LOC and orf annotations) with at least one FdNS mutation that appears specific to the lowland yak (Table S4). In addition, we used aggregated SIFT scores to estimate the predicted effect that the mutations had on each protein. Gene ontology enrichment analysis of the resulting complete gene list highlighted several biological processes including sperm axoneme assembly, G protein-coupled receptor signaling pathway, and perception of smell (Table S5).

*MUC20* was the only gene with a start-loss (start codon) high impact FdNS mutation, while other mucus protein genes *MUC3A* and *MUC16* (Fig. 3B) had one and five FdNS mutations, respectively, a signal we suggest could represent changes in the immune response (Montero et al. 2025). We found eight *KRT* genes presenting lowland yak-specific mutations with up to five FdNS mutations in one of them, *KRT38* (Fig. 3B), a gene that was found to have significant effect on wool traits in sheep (Sulayman et al. 2018). We identified *PLCB1*, *RNF26* and *DISP3*, which were previously linked to coat color in modern yaks (G. Li et al. 2022; X. Wu et al. 2024; Fu et al. 2026), and *SYNJ1*, part of the horn development region in yak (Liang et al. 2016). We also identified *AHNAK2*, *LYST, VPS13B* (three FdNS mutations) and *PCLO*, four genes that respectively relate to hair follicle, the Cohen syndrome characterized by thick hair, cattle coat colour, and body size (Momtazmanesh et al. 2020; Chovatiya et al. 2021; Reissmann and Ludwig 2013; Mukherjee et al. 2010), and all four genes have previously been highlighted in the cold-adapted woolly mammoth (Díez-del-Molino et al. 2023). Additionally, we report *ADGRB2*, previously suggested as a candidate gene for yak body size (Xinrui Liu et al. 2023).

We found several genes involved in stress response such as *MPV17L* involved in oxidative stress (Krick et al. 2008), *TMEM207*, *ISCA2* and *UQCRB*, also shown to interact with HIFs (Bunai et al. 2018; Green et al. 2022; Bao et al. 2021) as well as *COL15A1* and *POLQ*, which were previously linked to hypoxia and high-altitude adaptation in modern yaks (Qi et al. 2019; Yu et al. 2025). Additionally, *PPY*, *UCN2* and *QRFP* each produce neuropeptides involved in appetite control (Batterham et al. 2003; Inoue et al. 2003; Chartrel et al. 2003). We also identified a signal related to temperature and fat metabolism. *ADGRV1* and *CCDC168* (Fig. 3B), previously linked to cattle sweat glands and heat tolerance (Hernandez et al. 2024), had three and ten FdNS mutations, respectively. *DCLK3* (four FdNS mutations) was selected in a heat-resistant cattle breed (Xu et al. 2025), while *LOC616782* (six FdNS mutations; Fig. 3B) was highlighted in cattle heat-stress transcriptomics (Y. Liu et al. 2026). *CCDC57* (four FdNS mutations) was linked to milk fatty acid composition under heat stress in cattle (Bohlouli et al. 2022). *THADA* (three FdNS mutations) is involved in balancing energy metabolism and thermogenesis, selected in humans across different climates, and even associated with cold sensitivity in *Drosophila* knockouts (Moraru et al. 2017). *TET1* was already identified in woolly mammoths and may be connected to cold tolerance via adipocytes and repression of thermogenic genes such as *PPARGC1A*, which we also reported with one FdNS mutation in the lowland yak (Damal Villivalam et al. 2020; Díez-del-Molino et al. 2023). We also found *APOB*, a fat metabolism gene suggested to be under convergent evolution across Arctic mammals (Díez-del-Molino et al. 2023), while its associated receptor *APOBR* presented three FdNS mutations. We found one FdNS mutation in *TRPA1*, a gene highlighted in cold adaptation in both woolly mammoth and woolly rhinoceros (Lynch et al. 2015; Lord et al. 2020). Interestingly, *ARNTL2* lies at the crossroad between circadian rhythm, hypoxia and fat metabolism (Peek 2020; Mandl et al. 2022; G. Wu et al. 2023).

Finally, we specifically investigated the sequence in *EPAS1*, a gene encoding HIF-2α, selected in several taxa including hominins, horse or sheep, and associated with adaptation to the high-altitude hypoxic environment of the QTP (Huerta-Sánchez et al. 2014; Wei et al. 2016; Xuexue Liu et al. 2019). We show a shared derived state motif across exonic regions of the lowland, mountain and domestic yaks compared to other bovines not adapted to high-altitude (Fig. 3C). The overall clustering differs from the species tree while still grouping all lowland and mountain yaks together with a flat tree topology suggesting little variation in *EPAS1* exonic regions across yaks (Figs. 3C, S8-S9). The gain-of-function missense mutation H733Y fixed in wild mountain yaks and associated with increased *EPAS1* expression (Xinfeng Liu et al. 2023) was also identified in the four SnpEff-annotated lowland yak genomes. This shared *EPAS1* motif is possibly consistent with a pattern observed in snow leopards (L. Yang et al. 2025), in which the derived *EPAS1* allele was present in the Altai-Mongolian Plateau, and increased in frequency with altitude to near fixation on the QTP.

## Discussion

We generated the first genomic data from morphologically identified specimens of the Baikal yak and confirmed that it is the same taxon as the genomically assigned yak X (Fig. 1) (Gilardet et al. 2025; Oppenheimer, Gilardet, et al. 2026), which we now term the lowland yak (*Bos baikalensis*). In 1954, N. Verestchagin described the first partial skull holotype of the Baikal yak and tentatively attributed it to *Poephagus baikalensis* (Verestchagin 1954), although later investigation of an extensive collection of remains led to its reclassification as a subspecies: *Bos mutus baikalensis* (Vasiliev 2021). Following extensive palaeogenomics analyses demonstrating that yak X represents a distinct species to modern yaks (Oppenheimer, Gilardet, et al. 2026), and based on article 23 for the principle of priority from the International Code of Zoological Nomenclature (International Commission on Zoological Nomenclature 1999), we therefore revert the current palaeontological subspecies name back to the rank of species to yield the binomial: *Bos baikalensis* Verestchagin 1954 (Fig. 4). Based on its updated geographical range centered on the Altai-Sayan-Khangai Mountains, Transbaikalia and neighbouring steppe, and in comparison with the modern mountain yak located at much higher elevations on the QTP (Fig. 2), we propose to commonly refer to this extinct species as the lowland yak. Based on skull and horn morphology, the lowland yak was 12-15% larger than modern yaks, thus highlighting differentiation beyond DNA sequence data, while exhibiting pronounced sexual dimorphism similar to modern yaks (Vasiliev 2021).

**Figure 4:**
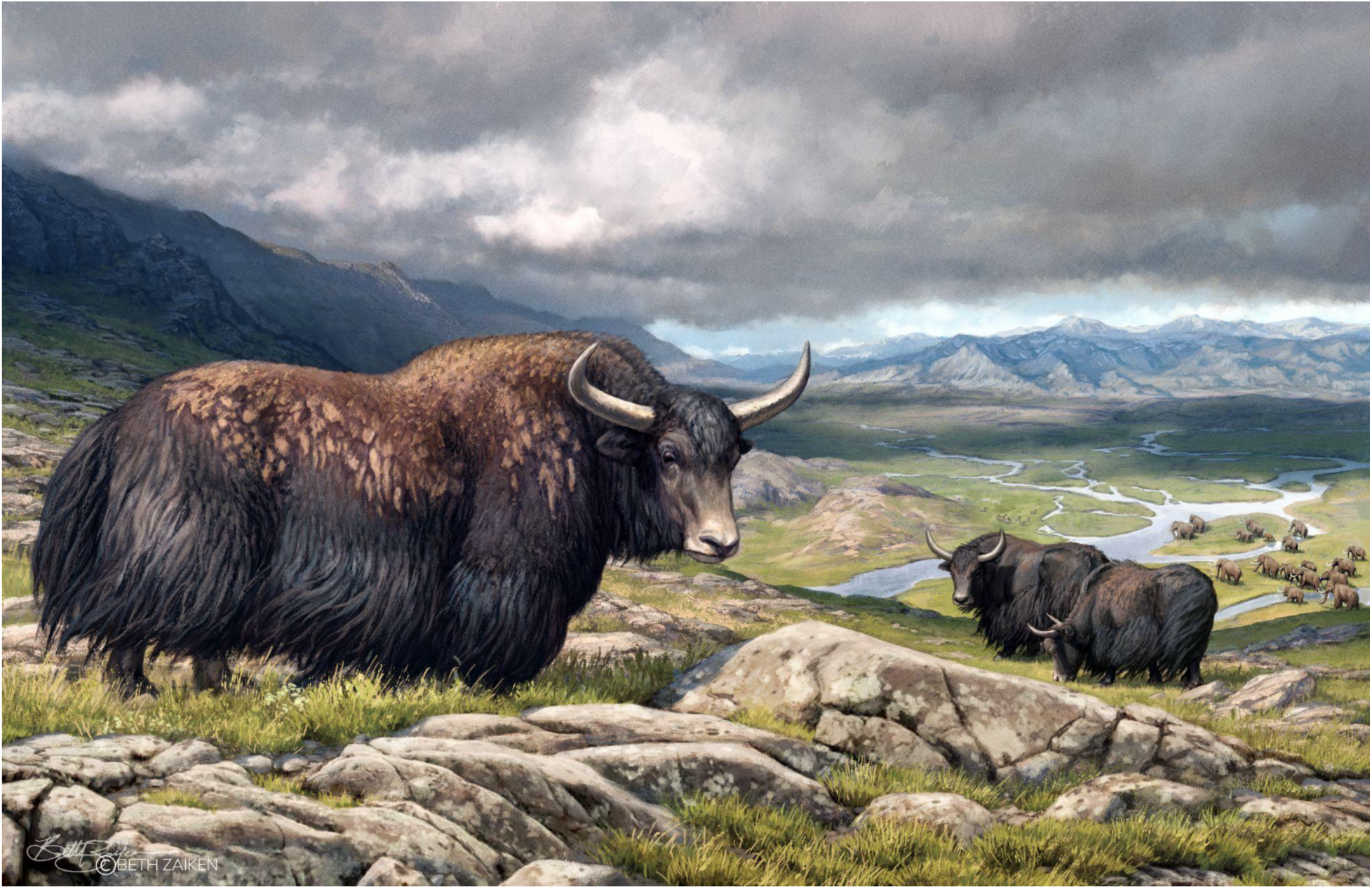
Illustration of the Pleistocene lowland yak by Beth Zaiken. made on the basis of the lowland yak’s larger size and the environment it inhabited (Vasiliev 2021).

Using multiple methods, we detected highly differentiated regions and genes in the lowland yak related to hair morphology, hypoxia, appetite control, fat metabolism and temperature. This signal is consistent with its environment ranging at lower altitude and characterized by better food availability compared to the QTP (Vasiliev 2021). Other signals in reproductive, immune and olfactory genes are consistent with these processes and genes being targets for rapid evolution across mammalian species (Torgerson et al. 2002; Kosiol et al. 2008; Niimura et al. 2014), and are reported in the evolution of other taxa such as horses (Orlando et al. 2013) and mammoths (Díez-del-Molino et al. 2023). However, we draw attention to potential false positives due to important gene length and mutational heterogeneity, as previously highlighted in olfactory receptors, *MUC16* and *PCLO* (M. S. Lawrence et al. 2013). Previous adaptation scans in modern yaks compared to cattle already highlighted hypoxia, energy metabolism, and sensory perception genes (Qiu et al. 2012). Finally, based on the highly similar coding sequence in the high-altitude-associated *EPAS1* gene observed between mountain and lowland yaks (Fig. 3C), we hypothesize that the lowland yak may have originated through a northward expansion from the QTP, where their common ancestor was adapted to the high elevations of the QTP. This would imply transient connectivity between the QTP and the range inhabited by the Pleistocene lowland yak (Herzschuh 2006; Cai et al. 2025).

Integrating morphological, palaeontological, geographical, and genomic information has provided an extended and refined understanding of the chronology of the extinct lowland yak (*Bos baikalensis*). The earliest yak remains are possibly from the Early Pleistocene in Kazakhstan (Flerow 1980), geographically located between the ranges of lowland and mountain yaks. However, this fossil predates the divergence of lowland and mountain yaks by at least 500,000 years (Oppenheimer, Gilardet, et al. 2026). The earliest evidence for lowland yak, derived from palaeogenomics, is from the Altai during Marine Isotope Stage (MIS) 7 of the Middle Pleistocene, before 200,000 years ago (Oppenheimer, Gilardet, et al. 2026). By the Late Pleistocene, the lowland yak inhabited the western side of its range (Altai and Khakassia) until the later stages of MIS 3 (Karginsky interstadial, ca. 50,000 - 22,000 years ago (Anderson and V. Lozhkin 2001)), and up to the beginning of the Last Glacial Maximum (LGM) (ca. 29,000 - 20,000 years ago, Sartanian MIS 2) in its eastern and southern ranges (Transbaikalia and Mongolia) (Vasiliev 2021). Although the timing of the lowland yak’s origins was previously estimated to MIS 11c (426,000 - 396,000 years ago (Tzedakis et al. 2022)) based on genomics (Oppenheimer, Gilardet, et al. 2026), future work should focus on elucidating the geographical origins of the lowland yak as well as the precise timing and location of its extinction.

## Materials and Methods

### Sample information

We generated genetic data from two specimens morphologically identified as Baikal yak (Vasiliev 2021) (Figs. 1A, S1; Table S1 and Supplementary Information). AG279 (collection ID: DCE-10) is a male metacarpal excavated at Denisova Cave (Fig. 2), from layer 11.3 of the East Chamber, which is dated to 80,000 ± 9,000 - 70,000 ± 8,000 years ago (Jacobs et al. 2019). AG281 is a female humerus excavated from the Fanatikov Cave (Fig. 2) and previously radiocarbon dated to >50,000 years ago (Vasiliev 2021).

### Sampling, DNA preparation and sequencing

Laboratory work was carried out in a dedicated ancient DNA facility. Two subsamples were taken from each specimen. A 43 and 96 mg subsample was cut off using a wire cutting tool, and an additional 20 and 31 mg of bone powder was drilled using an electric drill, from AG279 and AG281, respectively. Only bone fragments were pretreated in a 0.5%-0.7% sodium hypochlorite solution for 4 minutes, and subsequently rinsed three times in double-distilled water (Boessenkool et al. 2017). Each subsample was then lysed overnight and DNA was extracted following the Dabney and Meyer protocol (Dabney et al. 2013; Korlević et al. 2015). Double-stranded USER-treated libraries were generated following Meyer and Kircher (Meyer and Kircher 2010). Libraries were subsequently dual-indexed, and short fragments were removed using Cytiva beads with a 1.6× ratio. Purified libraries were assessed on a TapeStation with a D1000 kit prior to sequencing on an Illumina NovaSeq X platform using a 2×150 bp chemistry at the National Genomics Infrastructure (Science for Life Laboratory, Stockholm, Sweden).

### Data processing

We removed adapter sequences from the sequencing data and merged overlapping read pairs with SeqPrep2 (https://github.com/jeizenga/SeqPrep2), and retained only merged reads with a length of at least 35 bp. Subsequently, preprocessed reads were aligned to the ARS-UCD1.2 cattle (*Bos taurus*) genome, with a Y chromosome appended (from Btau5.1), using BWA v0.7.17-r1188 aln (H. Li and Durbin 2009) and the parameters -n 0.01 -o 2 -l 16500 (Prüfer et al. 2014). We then removed duplicates using Picard MarkDuplicates (https://broadinstitute.github.io/picard).

We determined genetic sex by comparing the number of reads (mapping quality MQ≥25) mapping to the X chromosome relative to autosomes, normalized by length (Shirazi et al. 2022; Oppenheimer, Gilardet, et al. 2026).

We called pseudohaploid genotypes at a set of 9.28 million transversion variants previously ascertained in a range of bovine species, including yak, gaur, banteng, cattle, steppe bison, European bison, and American bison (Oppenheimer, Gilardet, et al. 2026), using samtools (Danecek et al. 2021) mpileup, with the parameters -q20 -Q30 -B, and pileupCaller (https://github.com/stschiff/sequencetools). Pseudohaploid genotypes from AG279 and AG281 were then merged with pseudohaploid genotype calls on the same set of variants for 113 individuals from a range of bovine species, including yak X, domestic and wild mountain yak, bison, wisent, gaur, aurochs, taurine and zebu cattle, and water buffalo (Ji et al. 2021; Medugorac et al. 2017; Ahmad et al. 2025; N. Chen et al. 2020; Gelabert et al. 2021; Heaton et al. 2016; Lan et al. 2018; Qiu et al. 2015; Rossi et al. 2024; T. Sun et al. 2020; Shapiro et al. 2026; T. Yang et al. 2020; Verdugo et al. 2019; S. Zhang et al. 2021; D.-D. Wu et al. 2018; Węcek et al. 2017; Xinfeng Liu et al. 2023; Oppenheimer, Kapp, et al. 2026; Oppenheimer, Gilardet, et al. 2026) using mergeit (https://github.com/DReichLab/AdmixTools) (Table S6). This merged set of bovine pseudohaploid genotype calls was used for downstream *f*-statistic-based analyses.

We also mapped 32 publicly available modern bovine genomes (Oppenheimer et al. 2021; Xinfeng Liu et al. 2023; B. D. Rosen et al. 2020; K. Wang et al. 2017; Shapiro et al. 2026; T. Yang et al. 2020; Qiu et al. 2015; Heaton et al. 2016; Lan et al. 2018; Verdugo et al. 2019; T. Sun et al. 2020; D.-D. Wu et al. 2018) to the cattle reference for the *EPAS1* analysis, which required diploid genotype calls (Table S6). Adapter sequences were removed using Trimmomatic v0.39 (Bolger et al. 2014) with a minimum read length of 30 bp. Trimmed reads were subsequently aligned using BWA MEM (H. Li 2013), and duplicate reads were removed using Picard MarkDuplicates. We subsequently called diploid genotypes for these modern genomes using GATK (Auwera and O’Connor 2020).

We similarly generated diploid genotype calls from the four lowland yak individuals with >3-fold depth of coverage for both the FdNS and *EPAS1* analyses (Table S6). Diploid genotypes were called using the ancient DNA genotype caller snpAD (Prüfer 2018) with parameter -B and a minimum MQ of 25, following similar preprocessing, mapping and deduplication to the Baikal yak samples generated in this study.

Diploid genotypes were filtered for both depth and quality. Sites with a depth below ⅓× or above 3× of the mean depth were set as missing data using bcftools (Danecek et al. 2021) with the plugin +setGT, whereas a genotype quality minimum of 20 or 30 for ancient snpAD-called or modern GATK-called genotypes was applied, respectively. For the FdNS mutation analysis, we further restricted variants to only non-clustered SNPs using bcftools filter -g 5 and bcftools view-v snps, applied a mappability mask previously described (Oppenheimer, Gilardet, et al. 2026), and only retained variants overlapping autosomal exonic regions as annotated in the cattle reference genome.

### Placing Baikal yak within Bos and Bison diversity

We placed the mitochondrial and nuclear genomes of our two Baikal yak specimens into phylogenies to resolve their relationship to other bovine species, including yak X. First, for the mitochondrial phylogeny, we assembled mitogenomes from preprocessed reads using MIA (https://github.com/mpieva/mapping-iterative-assembler) on a wild mountain yak seed (NC_025563.1) using parameters -c -C -U -F -k 14 -s with an ancient DNA substitution matrix. We filtered consensus mitogenomes for a minimum of 3-fold depth of coverage and 66% agreement of the base called (https://github.com/aersoares81/mia-helper-scripts). We aligned our two filtered mitogenomes using MUSCLE v.3.8.31 (Edgar 2004) to a set of 76 mitogenomes including yak X, modern yak, and bison (Oppenheimer, Gilardet, et al. 2026), and subsequently filtered sites for a maximum of 20% missingness (Swali et al. 2023). We generated the tree using IQ-TREE v.2.4.0 -m MFP -bb 1000 -alrt 1000 (Nguyen et al. 2015; Kalyaanamoorthy et al. 2017) with a best-fitted HKY + F + I model. The tree was subsequently visualised using the Interactive Tree of Life (Letunic and Bork 2024).

For the autosomal phylogeny, we calculated all pairwise *f2*-statistics from the merged pseudohaploid calls using fstats.py (https://github.com/jooppenh/fstats) (Oppenheimer, Gilardet, et al. 2026). These were used to construct a neighbor-joining tree using ape (Paradis and Schliep 2019), which was rooted using water buffalo, and visualized using the Interactive Tree of Life. Branch support was generated via weighted blocked bootstrapping using 5 Mb windows and 1000 replicates.

We additionally constructed a PCA using a matrix of all possible combinations of *f3*-statistics of the form *f3*(X, A, B), where X, A, and B are different individual yaks, including modern domestic and wild mountain yaks, yak X and the two individuals morphologically identified as Baikal yaks. All *f3*-statistics were calculated using fstats.py and the PCA was generated using prcomp in R (https://www.rdocumentation.org/packages/stats/versions/3.6.2/topics/prcomp).

### Allele sharing tests

Allele sharing statistics were used to evaluate whether the Baikal yak formed a clade with yak X, or instead shared ancestry with modern yak, bison and/or aurochs. All D-statistics of the form D(yak X, Baikal yak; wild mountain yak, water buffalo), D(yak X, Baikal yak; steppe bison, American bison) or D(yak X, Baikal yak; Asian aurochs, European aurochs) were calculated using admixtools2 (Maier et al. 2023) with auto_only=FALSE and f4mode=FALSE, and filtered for at least 100,000 SNPs within any individual statistic. We also tested the affinity of the Baikal yak specimens to other yaks by calculating D(yak X, wild mountain yak; Baikal yak, water buffalo) in order to confirm that Baikal yaks have excess allele sharing with yak X, relative to other yaks, instead of representing a separate lineage equally related to both wild mountain yak and yak X.

Since Mongolian yaks are the modern population which is geographically closest to the past range of lowland yaks, we additionally specifically tested for lowland yak ancestry in Mongolian domestic yaks relative to other domestic yaks by calculating D(Mongolian domestic yak, Chinese domestic yak; lowland yak, bison), and tested for allele sharing between Mongolian and Chinese domestic yaks, relative to lowland yaks, by calculating D(Mongolian domestic yak, lowland yak; Chinese domestic yak, bison).

### Functional changes analyses

To identify highly differentiated genomic regions in the lowland yak, we calculated a Population Branch Statistic (PBS) with *f3*-statistics to test for excessively long branches in lowland yak relative to other yaks and bison. For this, we grouped individuals by species (Table S6), using lowland yak genomes with depth of coverage >0.2-fold, including AG279 and AG281, and calculated *f3*(lowland yak; wild mountain yak, American bison) in sliding 500 kb windows with a 10 kb step using fstats.py, requiring a minimum of 200 and maximum of 10,000 SNPs in each window. Per-window *f3-*statistic values were converted to *Z*-scores by randomly selecting 388 windows at least 5 Mb apart and calculating their mean and standard deviation. We tested different outgroup species used for computing the PBS, also calculating *f3*(lowland yak; domestic yak, American bison) and *f3*(lowland yak; wild mountain yak, domestic yak) with similar random window selection. To assess peaks and overlapping genes, we subsequently intersected genomic regions containing windows with *Z*-scores above 5 and at least 950 SNPs with annotations in the cattle reference genome.

We investigated possible fixed-derived non-synonymous (FdNS) mutations in lowland yak. We identified putative FdNS variants in the four lowland yak genomes with depth of coverage >3-fold using SnpEff version 4.3t (Cingolani et al. 2012) and the cattle annotations, and subsequently called the same sites using bcftools for comparison in a reference panel of four wild mountain yaks and four bison, with a minimum MQ of 25 (Table S6). To classify a variant as fixed, we applied maximum missingness and Minor Allele Frequency (MAF) filters of 0.25 and 0.025, respectively. We then computed the SIFT score for effect prediction of the remaining variants (Ng and Henikoff 2003). The aggregated SIFT score we present is the sum of (1 - SIFT score) for every variant in a given gene. We normalized the number of FdNS mutations by gene length, which was obtained using the agat_sp_keep_longest_isoform.pl script (https://github.com/NBISweden/AGAT) to select the longest isoform, and GffRead (Pertea and Pertea 2020). The pipeline for identifying putative FdNS mutations is available at https://github.com/NathanACO/Selection_Pipeline. Gene ontology enrichment analysis for biological processes (complete annotation dataset) was performed using PANTHER (https://geneontology.org/) with *Bos taurus* as reference and default parameters.

We directly investigated the high-altitude-related gene *EPAS1* on a set of 36 individuals across bovine diversity, including the four lowland yak >3-fold depth of coverage, wild mountain and domestic yaks, American bison, wisent, gaur, banteng, taurine and zebu cattle, and water buffalo (Table S6). After merging individuals, we extracted *EPAS1* exons as annotated in the reference genome, and plotted sites in R using packages vcfR (Knaus and Grünwald 2017) and pheatmap (https://pheatmap.com/), while setting multiallelic sites as missing data. We additionally generated a maximum-likelihood phylogeny by converting the VCF to phylip format using vcf2phylip.py (https://github.com/edgardomortiz/vcf2phylip) and using IQ-TREE v2.4 with parameters -nt AUTO -m MFP -bb 1000 -alrt 1000. We also generated an alternative tree version covering the whole *EPAS1* gene (chr11:28,735,330-28,825,665), after filtering sites for a maximum of 20% missingness. Trees were visualized in FigTree (https://github.com/rambaut/figtree).

## Data availability

Sequencing data for voucher specimens AG279 and AG281 are available on ENA project PRJEB129787.

## Author contributions

Conceptualization: AG, JO, PDH, LD

Funding acquisition: PDH, LD

Resources: AG, MBK, EE, MVS, SKV

Investigation: AG, MBK, GX, SKV, KD

Data curation: AG, JO

Formal analysis: AG, JO, NM

Visualization: AG, JO, MBK

Writing – original draft: AG

Writing – review & editing: all authors

## Acknowledgements

We thank Elena Essel for her valuable logistical help, and Beth Zaiken for her illustration of the lowland yak. This study was funded by the Swedish Research Council (VR grant 2021-00625), the European Union (ERC grant ‘PrimiGenomes’ 101054984), and the Knut and Alice Wallenberg Foundation (KAW grants 2022.0033 and 2021.0048). Data analysis was supported by the National Academic Infrastructure for Supercomputing in Sweden (NAISS) (projects 2026/1-30 and 2025/22-1635). Archeological work at Denisova Cave was funded by the Russian Science Foundation (grant 24-78-10006), and work on the morphology and primary taxonomy of the materials at Fanatikov Cave was associated with project FWZG-2026-0002. N.M. gratefully acknowledges support from the Human Frontier Science Program (HFSP; RGP023/2024)

## Notes

**Conflicts of interest:** The authors declare no conflict of interest.

### Competing Interest Statement

The authors have declared no competing interest.

## References

Ahmad, Sheikh Firdous, Munish Gangwar, Amit Kumar, et al. 2025. “Dissecting Genomes of Multiple Yak Populations: Unveiling Ancestry and High-Altitude Adaptation through Whole-Genome Resequencing Analysis.” BMC Genomics 26 (1): 214.

Anderson, Patricia M., and Anatoly V. Lozhkin. 2001. “The Stage 3 Interstadial Complex (Karginskii/Middle Wisconsinan Interval) of Beringia: Variations in Paleoenvironments and Implications for Paleoclimatic Interpretations.” Quaternary Science Reviews 20 (1–3): 93–125.

Auwera, Geraldine van der, and Brian D. O’Connor. 2020. Genomics in the Cloud: Using Docker, GATK, and WDL in Terra. O’Reilly Media, Incorporated.

Bao, Xingting, Jinhua Zhang, Guomin Huang, et al. 2021. “The Crosstalk between HIFs and Mitochondrial Dysfunctions in Cancer Development.” Cell Death & Disease 12 (2): 215.

Baquero, Monica M., and Brandon L. Plattner. 2017. “Bovine Peripheral Blood WC1+ and WC1neg Γδ T Lymphocytes Modulate Monocyte-Derived Macrophage Effector Functions during in Vitro Mycobacterium Avium Subspecies Paratuberculosis Infection.” Cellular Immunology 315 (May): 34–44.

Batterham, R. L., C. W. Le Roux, M. A. Cohen, et al. 2003. “Pancreatic Polypeptide Reduces Appetite and Food Intake in Humans.” The Journal of Clinical Endocrinology and Metabolism 88 (8): 3989–92.

Baum, D. A., and K. L. Shaw. 1995. “Genealogical Perspectives on the Species Problem.” In Experimental and Molecular Approaches to Plant Biosystematics, edited by P. C. Hoch and A. G. Stephenson.

Boessenkool, Sanne, Kristian Hanghøj, Heidi M. Nistelberger, et al. 2017. “Combining Bleach and Mild Predigestion Improves Ancient DNA Recovery from Bones.” Molecular Ecology Resources 17 (4): 742–51.

Bohlouli, M., K. Halli, T. Yin, N. Gengler, and S. König. 2022. “Genome-Wide Associations for Heat Stress Response Suggest Potential Candidate Genes Underlying Milk Fatty Acid Composition in Dairy Cattle.” Journal of Dairy Science 105 (4): 3323–40.

Bolger, Anthony M., Marc Lohse, and Bjoern Usadel. 2014. “Trimmomatic: A Flexible Trimmer for Illumina Sequence Data.” Bioinformatics (Oxford, England) 30 (15): 2114–20.

Bunai, Katsuaki, Hiroshi Okubo, Kimika Hano, et al. 2018. “TMEM207 Hinders the Tumour Suppressor Function of WWOX in Oral Squamous Cell Carcinoma.” Journal of Cellular and Molecular Medicine 22 (2): 1026–33.

Burrell, Andrew S., Todd R. Disotell, and Christina M. Bergey. 2015. “The Use of Museum Specimens with High-Throughput DNA Sequencers.” Journal of Human Evolution 79 (February): 35–44.

Buzzard, Paul J., Donghua Xu, and Huan Li. 2014. “Sexual/Aggressive Behavior of Wild Yak (Bos Mutus Prejevalsky 1883) during the Rut: Influence of Female Choice.” Kexue Tongbao [Chinese Science Bulletin] 59 (22): 2756–63.

Cai, Jing, Gang Liu, and Jianquan Liu. 2025. “Biogeographic Connections between the Flora of the Arctic and Qinghai-Tibet Plateau: Building on the Legacy of Richard Abbott.” Plant Ecology & Diversity 18 (3–4): 87–98.

Chartrel, Nicolas, Cynthia Dujardin, Youssef Anouar, et al. 2003. “Identification of 26RFa, a Hypothalamic Neuropeptide of the RFamide Peptide Family with Orexigenic Activity.” Proceedings of the National Academy of Sciences of the United States of America 100 (25): 15247–52.

Chen, Mengge, Xuancheng Zhang, Xiaocen Wang, et al. 2026. “GBP5-Triggered AIM2 Inflammasome Drives Host Defense and Exacerbates Disease Severity during Neospora Caninum Infection.” Veterinary Research 57 (1): 113.

Chen, Ningbo, Lele Ren, Linyao Du, et al. 2020. “Ancient Genomes Reveal Tropical Bovid Species in the Tibetan Plateau Contributed to the Prevalence of Hunting Game until the Late Neolithic.” Proceedings of the National Academy of Sciences of the United States of America 117 (45): 28150–59.

Chiou, Kenneth L., Mareike C. Janiak, India A. Schneider-Crease, et al. 2022. “Genomic Signatures of High-Altitude Adaptation and Chromosomal Polymorphism in Geladas.” Nature Ecology & Evolution 6 (5): 630–43.

Chovatiya, Gopal, Sangeeta Ghuwalewala, Lauren D. Walter, Benjamin D. Cosgrove, and Tudorita Tumbar. 2021. “High-Resolution Single-Cell Transcriptomics Reveals Heterogeneity of Self-Renewing Hair Follicle Stem Cells.” Experimental Dermatology 30 (4): 457–71.

Cingolani, Pablo, Adrian Platts, Le Lily Wang, et al. 2012. “A Program for Annotating and Predicting the Effects of Single Nucleotide Polymorphisms, SnpEff: SNPs in the Genome of Drosophila Melanogaster Strain W1118; Iso-2; Iso-3: SNPs in the Genome of Drosophila Melanogaster Strain w ^1118^; Iso-2; Iso-3.” Fly 6 (2): 80–92.

Dabney, Jesse, Michael Knapp, Isabelle Glocke, et al. 2013. “Complete Mitochondrial Genome Sequence of a Middle Pleistocene Cave Bear Reconstructed from Ultrashort DNA Fragments.” Proceedings of the National Academy of Sciences of the United States of America 110 (39): 15758–63.

Damal Villivalam, Sneha, Dongjoo You, Jinse Kim, et al. 2020. “TET1 Is a Beige Adipocyte-Selective Epigenetic Suppressor of Thermogenesis.” Nature Communications 11 (1): 4313.

Damgaard, Peter B., Ashot Margaryan, Hannes Schroeder, Ludovic Orlando, Eske Willerslev, and Morten E. Allentoft. 2015. “Improving Access to Endogenous DNA in Ancient Bones and Teeth.” Scientific Reports 5 (1): 11184.

Danecek, Petr, James K. Bonfield, Jennifer Liddle, et al. 2021. “Twelve Years of SAMtools and BCFtools.” GigaScience 10 (2): giab008.

Díez-del-Molino, David, Marianne Dehasque, J. Camilo Chacón-Duque, et al. 2023. “Genomics of Adaptive Evolution in the Woolly Mammoth.” Current Biology 33 (9): 1753–1764.e4.

Donoghue, Michael J. 1985. “A Critique of the Biological Species Concept and Recommendations for a Phylogenetic Alternative.” The Bryologist 88 (3): 172–81.

Edgar, Robert C. 2004. “MUSCLE: Multiple Sequence Alignment with High Accuracy and High Throughput.” Nucleic Acids Research 32 (5): 1792–97.

Eisenmann, Véra, and Sergej Vasiliev. 2011. “Unexpected Finding of a New Equus Species (Mammalia, Perissodactyla) Belonging to a Supposedly Extinct Subgenus in Late Pleistocene Deposits of Khakassia (Southwestern Siberia).” Geodiversitas 33 (3): 519–30.

Flerow, Constantin C. 1980. “Zur geographischen Verbreitung der Gattung Poephagus im Pleistozän und Holozän.” In Band 4, edited by H. D. Kahlke. De Gruyter.

Fu, Yujiao, Jiahong Zhao, Yuanyuan Yu, et al. 2026. “Genome-Wide Detection and Genomic Selection Based on Genotype-by-Environment Interaction-Associated Signals for Yak Coat Color.” BMC Genomics 27 (1): 805.

Gelabert, Pere, Susanna Sawyer, Anders Bergström, et al. 2021. “Genome-Scale Sequencing and Analysis of Human, Wolf, and Bison DNA from 25,000-Year-Old Sediment.” Current Biology 31 (16): 3564–3574.e9.

Gilardet, Alexandre, Jonas Oppenheimer, Mikkel-Holger S. Sinding, et al. 2025. “Paleogenomics Reveals a Loss of Bovine Lineages in Mid-Latitude Asia Over the Last 200,000 Years.” Genome Biology and Evolution 17 (11): evaf206.

Green, Yangsook Song, Maria C. Ferreira dos Santos, Daniel G. Fuja, et al. 2022. “ISCA2 Inhibition Decreases HIF and Induces Ferroptosis in Clear Cell Renal Carcinoma.” Oncogene 41 (42): 4709–23.

Hao, Dan, Xiao Wang, Bo Thomsen, et al. 2020. “Copy Number Variations and Expression Levels of Guanylate-Binding Protein 6 Gene Associated with Growth Traits of Chinese Cattle.” Animals: An Open Access Journal From MDPI 10 (4): E566.

Heaton, Michael P., Timothy P. L. Smith, Jacky K. Carnahan, et al. 2016. “Using Diverse U.S. Beef Cattle Genomes to Identify Missense Mutations in EPAS1, a Gene Associated with Pulmonary Hypertension.” F1000Research 5: 2003.

Heiner, Michael, Davaa Galbadrakh, and Joseph Kiesecker. 2019. “Shifting Winds in the Mongolian Gobi Desert: Nature and Traditions Face the Modern Era.” In Reference Module in Earth Systems and Environmental Sciences. Elsevier.

Heintzman, Peter D., Grant D. Zazula, Ross D. E. MacPhee, et al. 2017. “A New Genus of Horse from Pleistocene North America.” eLife 6 (November): e29944.

Hernandez, Aakilah S., Gabriel A. Zayas, Eduardo E. Rodriguez, et al. 2024. “Exploring the Genetic Control of Sweat Gland Characteristics in Beef Cattle for Enhanced Heat Tolerance.” Journal of Animal Science and Biotechnology 15 (1): 66.

Herzschuh, Ulrike. 2006. “Palaeo-Moisture Evolution in Monsoonal Central Asia during the Last 50,000 Years.” Quaternary Science Reviews 25 (1–2): 163–78.

Huerta-Sánchez, Emilia, Xin Jin, Asan, et al. 2014. “Altitude Adaptation in Tibetans Caused by Introgression of Denisovan-like DNA.” Nature 512 (7513): 194–97.

Inoue, Koki, Glenn R. Valdez, Teresa M. Reyes, et al. 2003. “Human Urocortin II, a Selective Agonist for the Type 2 Corticotropin-Releasing Factor Receptor, Decreases Feeding and Drinking in the Rat.” The Journal of Pharmacology and Experimental Therapeutics 305 (1): 385–93.

International Commission on Zoological Nomenclature. 1999. International Code of Zoological Nomenclature = Code International de Nomenclature Zoologique. 4th ed. Vol. 1999. International Trust for Zoological Nomenclature, c/o Natural History Museum.

Jacobs, Zenobia, Bo Li, Michael V. Shunkov, et al. 2019. “Timing of Archaic Hominin Occupation of Denisova Cave in Southern Siberia.” Nature 565 (7741): 594–99.

Jacobson-Tepfer, Esther. 2025. “Wild Yaks, Domesticated Yaks, and the Emergence of Transhumant Pastoralism in the Mongolian Altai.” Archaeological Research in Asia 43 (September): 100642.

Ji, Qiu-Mei, Jin-Wei Xin, Zhi-Xin Chai, et al. 2021. “A Chromosome-Scale Reference Genome and Genome-Wide Genetic Variations Elucidate Adaptation in Yak.” Molecular Ecology Resources 21 (1): 201–11.

Kalyaanamoorthy, Subha, Bui Quang Minh, Thomas K. F. Wong, Arndt von Haeseler, and Lars S. Jermiin. 2017. “ModelFinder: Fast Model Selection for Accurate Phylogenetic Estimates.” Nature Methods 14 (6): 587–89.

Knaus, Brian J., and Niklaus J. Grünwald. 2017. “Vcfr: A Package to Manipulate and Visualize Variant Call Format Data in R.” Molecular Ecology Resources 17 (1): 44–53.

Korlević, Petra, Tobias Gerber, Marie-Theres Gansauge, et al. 2015. “Reducing Microbial and Human Contamination in DNA Extractions from Ancient Bones and Teeth.” BioTechniques 59 (2): 87–93.

Kosiol, Carolin, Tomás Vinar, Rute R. da Fonseca, et al. 2008. “Patterns of Positive Selection in Six Mammalian Genomes.” PLoS Genetics 4 (8): e1000144.

Krick, Stefanie, Shaolin Shi, Wenjun Ju, et al. 2008. “Mpv17l Protects against Mitochondrial Oxidative Stress and Apoptosis by Activation of Omi/HtrA2 Protease.” Proceedings of the National Academy of Sciences of the United States of America 105 (37): 14106–11.

Lacroix-Pépin, Nicolas, Ghislain Danyod, Narayanan Krishnaswamy, et al. 2011. “The Multidrug Resistance-Associated Protein 4 (MRP4) Appears as a Functional Carrier of Prostaglandins Regulated by Oxytocin in the Bovine Endometrium.” Endocrinology 152 (12): 4993–5004.

Lan, Daoliang, Xianrong Xiong, Tserang-Donko Mipam, et al. 2018. “Genetic Diversity, Molecular Phylogeny, and Selection Evidence of Jinchuan Yak Revealed by Whole-Genome Resequencing.” G3 (Bethesda, Md.) 8 (3): 945–52.

Lawrence, Macy T., Trent E. Schwartz, Loni W. Lucherk, Ty E. Lawrence, and Emilie C. Baker. 2026. “Differential Gene Expression in Glisson’s Capsule Associated with Liver Abscesses in Feedlot Cattle.” Journal of Animal Science 104 (skag320): skag320.

Lawrence, Michael S., Petar Stojanov, Paz Polak, et al. 2013. “Mutational Heterogeneity in Cancer and the Search for New Cancer-Associated Genes.” Nature 499 (7457): 214–18.

Lebedev, Vladimir S., Yulia Kovalskaya, Evgeniya N. Solovyeva, et al. 2021. “Molecular Systematics of the Sicista Tianschanica Species Complex: A Contribution from Historical DNA Analysis.” PeerJ 9 (January): e10759.

Leonardi, Michela, Pablo Librado, Clio Der Sarkissian, et al. 2017. “Evolutionary Patterns and Processes: Lessons from Ancient DNA.” Systematic Biology 66 (1): e1–29.

Leslie, David M., Jr, and George B. Schaller. 2009. “Bos Grunniens and Bos Mutus (Artiodactyla: Bovidae).” Mammalian Species, no. 836 (May): 1–17.

Letunic, Ivica, and Peer Bork. 2024. “Interactive Tree of Life (iTOL) v6: Recent Updates to the Phylogenetic Tree Display and Annotation Tool.” Nucleic Acids Research 52 (W1): W78–82.

Li, Cai, Gerald Wiener, Gerald Wiener, Han Jianlin, and Long Ruijun. 2003. THE YAK - SECOND EDITION. Food and Agriculture Organization of the United Nations, Regional Office for Asia and the Pacific.

Li, Guangzhen, Jing Luo, Fuwen Wang, et al. 2022. “Whole-Genome Resequencing Reveals Genetic Diversity, Differentiation, and Selection Signatures of Yak Breeds/Populations in Qinghai, China.” Frontiers in Genetics 13: 1034094.

Li, Heng. 2013. “Aligning Sequence Reads, Clone Sequences and Assembly Contigs with BWA-MEM.” In arXiv [q-Bio.GN]. March 16. arXiv. 10.48550/arXiv.1303.3997.

Li, Heng, and Richard Durbin. 2009. “Fast and Accurate Short Read Alignment with Burrows–Wheeler Transform.” Bioinformatics (Oxford, England) 25 (14): 1754–60.

Liang, Chunnian, Lizhong Wang, Xiaoyun Wu, et al. 2016. “Genome-Wide Association Study Identifies Loci for the Polled Phenotype in Yak.” PloS One 11 (7): e0158642.

Liu, Xinfeng, Wenyu Liu, Johannes A. Lenstra, et al. 2023. “Evolutionary Origin of Genomic Structural Variations in Domestic Yaks.” Nature Communications 14 (1): 5617.

Liu, Xinrui, Mingxiu Wang, Jie Qin, et al. 2023. “Identification of Candidate Genes Associated with Yak Body Size Using a Genome-Wide Association Study and Multiple Populations of Information.” Animals: An Open Access Journal From MDPI 13 (9): 1470.

Liu, Xuexue, Yanli Zhang, Yefang Li, et al. 2019. “EPAS1 Gain-of-Function Mutation Contributes to High-Altitude Adaptation in Tibetan Horses.” Molecular Biology and Evolution 36 (11): 2591–603.

Liu, Yongjia, Zitai Guo, Na Cao, et al. 2026. “Coated Cobalamin Alleviates Heat Stress in Lactating Holstein Cows by Enhancing Ruminal Propionate Production, Antioxidant Capacity, and Hepatic Mitogen-Activated Protein Kinase Signaling.” Journal of Animal Science and Biotechnology 17 (1): 168.

Lord, Edana, Nicolas Dussex, Marcin Kierczak, et al. 2020. “Pre-Extinction Demographic Stability and Genomic Signatures of Adaptation in the Woolly Rhinoceros.” Current Biology 30 (19): 3871–3879.e7.

Lu, Huayu, Xianyan Wang, Xiaoyong Wang, et al. 2019. “Formation and Evolution of Gobi Desert in Central and Eastern Asia.” Earth-Science Reviews 194 (July): 251–63.

Lynch, Vincent J., Oscar C. Bedoya-Reina, Aakrosh Ratan, et al. 2015. “Elephantid Genomes Reveal the Molecular Bases of Woolly Mammoth Adaptations to the Arctic.” Cell Reports 12 (2): 217–28.

Mahmut, Halik, Ryuichi Masuda, Manabu Onuma, et al. 2002. “Molecular Phylogeography of the Red Deer (Cervus Elaphus) Populations in Xinjiang of China: Comparison with Other Asian, European, and North American Populations.” Zoological Science 19 (4): 485–95.

Maier, Robert, Pavel Flegontov, Olga Flegontova, Ulaş Işıldak, Piya Changmai, and David Reich. 2023. “On the Limits of Fitting Complex Models of Population History to F-Statistics.” eLife 12 (e85492): e85492.

Mandl, Markus, Hans P. Viertler, Maria Zopoglou, et al. 2022. “The Circadian Transcription Factor ARNTL2 Is Regulated by Weight-Loss Interventions in Human White Adipose Tissue and Inhibits Adipogenesis.” Cell Death Discovery 8 (1): 443.

Medugorac, Ivica, Alexander Graf, Cécile Grohs, et al. 2017. “Whole-Genome Analysis of Introgressive Hybridization and Characterization of the Bovine Legacy of Mongolian Yaks.” Nature Genetics 49 (3): 470–75.

Meyer, Matthias, and Martin Kircher. 2010. “Illumina Sequencing Library Preparation for Highly Multiplexed Target Capture and Sequencing.” Cold Spring Harbor Protocols 2010 (6): db.prot5448.

Momtazmanesh, Sara, Elham Rayzan, Sepideh Shahkarami, Meino Rohlfs, Christoph Klein, and Nima Rezaei. 2020. “A Novel VPS13B Mutation in Cohen Syndrome: A Case Report and Review of Literature.” BMC Medical Genetics 21 (June): 140.

Montero, Paula, Inés Roger, Javier Milara, and Julio Cortijo. 2025. “Immunomodulatory Properties of Transmembrane Mucins: From Chronic Diseases to Cancer.” Physiological Reviews 105 (4): 2233–304.

Moraru, Alexandra, Gulcin Cakan-Akdogan, Katrin Strassburger, et al. 2017. “THADA Regulates the Organismal Balance between Energy Storage and Heat Production.” Developmental Cell 41 (1): 72–81.e6.

Mukherjee, Konark, Xiaofei Yang, Stefan H. Gerber, et al. 2010. “Piccolo and Bassoon Maintain Synaptic Vesicle Clustering without Directly Participating in Vesicle Exocytosis.” Proceedings of the National Academy of Sciences 107 (14): 6504–9.

Ng, Pauline C., and Steven Henikoff. 2003. “SIFT: Predicting Amino Acid Changes That Affect Protein Function.” Nucleic Acids Research 31 (13): 3812–14.

Nguyen, Lam-Tung, Heiko A. Schmidt, Arndt von Haeseler, and Bui Quang Minh. 2015. “IQ-TREE: A Fast and Effective Stochastic Algorithm for Estimating Maximum-Likelihood Phylogenies.” Molecular Biology and Evolution 32 (1): 268–74.

Niimura, Yoshihito, Atsushi Matsui, and Kazushige Touhara. 2014. “Extreme Expansion of the Olfactory Receptor Gene Repertoire in African Elephants and Evolutionary Dynamics of Orthologous Gene Groups in 13 Placental Mammals.” Genome Research 24 (9): 1485–96.

Oppenheimer, Jonas, Alexandre Gilardet, Maxim B. Kozlikin, et al. 2026. “Ancient Genomes Reveal an Extinct Bovine Species from Mid-Latitude Asia.” Current Biology, ahead of print, October 5. 10.1016/j.cub.2026.09.043.

Oppenheimer, Jonas, Joshua D. Kapp, Molly Cassatt-Johnstone, et al. 2026. “Paleogenomic Insight into the Collapse, Recovery, and Management of American Bison.” Science (New York, N.Y.) 393 (6811). 10.1126/science.aee4205.

Oppenheimer, Jonas, Benjamin D. Rosen, Michael P. Heaton, et al. 2021. “A Reference Genome Assembly of American Bison, Bison Bison Bison.” The Journal of Heredity 112 (2): 174–83.

Orlando, Ludovic, Aurélien Ginolhac, Guojie Zhang, et al. 2013. “Recalibrating Equus Evolution Using the Genome Sequence of an Early Middle Pleistocene Horse.” Nature 499 (7456): 74–78.

Paradis, Emmanuel, and Klaus Schliep. 2019. “Ape 5.0: An Environment for Modern Phylogenetics and Evolutionary Analyses in R.” Bioinformatics (Oxford, England) 35 (3): 526–28.

Peek, Clara B. 2020. “Metabolic Implications of Circadian-HIF Crosstalk.” Trends in Endocrinology and Metabolism 31 (6): 459–68.

Pertea, Geo, and Mihaela Pertea. 2020. “GFF Utilities: GffRead and GffCompare.” F1000Research 9 (April): 304.

Prüfer, Kay. 2018. “snpAD: An Ancient DNA Genotype Caller.” Bioinformatics (Oxford, England) 34 (24): 4165–71.

Prüfer, Kay, Fernando Racimo, Nick Patterson, et al. 2014. “The Complete Genome Sequence of a Neanderthal from the Altai Mountains.” Nature 505 (7481): 43–49.

Qi, Xuebin, Qu Zhang, Yaoxi He, et al. 2019. “The Transcriptomic Landscape of Yaks Reveals Molecular Pathways for High Altitude Adaptation.” Genome Biology and Evolution 11 (1): 72–85.

Qiu, Qiang, Lizhong Wang, Kun Wang, et al. 2015. “Yak Whole-Genome Resequencing Reveals Domestication Signatures and Prehistoric Population Expansions.” Nature Communications 6 (1): 10283.

Qiu, Qiang, Guojie Zhang, Tao Ma, et al. 2012. “The Yak Genome and Adaptation to Life at High Altitude.” Nature Genetics 44 (8): 946–49.

Queiroz, Kevin de. 2005. “Ernst Mayr and the Modern Concept of Species.” Proceedings of the National Academy of Sciences 102 (suppl_1): 6600–6607.

Reich, David, Richard E. Green, Martin Kircher, et al. 2010. “Genetic History of an Archaic Hominin Group from Denisova Cave in Siberia.” Nature 468 (7327): 1053–60.

Reissmann, Monika, and Arne Ludwig. 2013. “Pleiotropic Effects of Coat Colour-Associated Mutations in Humans, Mice and Other Mammals.” Seminars in Cell & Developmental Biology 24 (6): 576–86.

Rosen, Benjamin D., Derek M. Bickhart, Robert D. Schnabel, et al. 2020. “De Novo Assembly of the Cattle Reference Genome with Single-Molecule Sequencing.” GigaScience 9 (3): giaa021.

Rosen, D. E. 1979. “Fishes from the Uplands and Intermontane Basins of Guatemala : Revisionary Studies and Comparative Geography.” Bulletin of the American Museum of Natural History 162: 267–376.

Rossi, Conor, Mikkel-Holger S. Sinding, Victoria E. Mullin, et al. 2024. “The Genomic Natural History of the Aurochs.” Nature 635 (8037): 136–41.

Schaller, George B. 1998. Wildlife of the Tibetan Steppe. University of Chicago Press.

Shapiro, Beth, Jonas Oppenheimer, Michael P. Heaton, et al. 2026. “Most Beefalo Cattle Have No Detectable Bison Genetic Ancestry.” eLife 13 (June). 10.7554/eLife.102750.2.

Shirazi, Sabrina, Nasreen Broomandkhoshbacht, Jonas Oppenheimer, et al. 2022. “Ancient DNA-Based Sex Determination of Bison Hide Moccasins Indicates Promontory Cave Occupants Selected Female Hides for Footwear.” Journal of Archaeological Science 137 (January): 105533.

Simpson, George Gaylord. 1961. Principles of Animal Taxonomy. Columbia University Press.

Sokal, Robert R., and Theodore J. Crovello. 1970. “The Biological Species Concept: A Critical Evaluation.” The American Naturalist 104 (936): 127–53.

Sulayman, Ablat, Mahira Tursun, Yiming Sulaiman, et al. 2018. “Association Analysis of Polymorphisms in Six Keratin Genes with Wool Traits in Sheep.” Asian-Australasian Journal of Animal Sciences 31 (6): 775–83.

Sun, Jimin, Weiguo Liu, Zhonghui Liu, Tao Deng, Brian F. Windley, and Bihong Fu. 2017. “Extreme Aridification since the Beginning of the Pliocene in the Tarim Basin, Western China.” Palaeogeography, Palaeoclimatology, Palaeoecology 485 (November): 189–200.

Sun, Ting, Jiafei Shen, Alessandro Achilli, et al. 2020. “Genomic Analyses Reveal Distinct Genetic Architectures and Selective Pressures in Buffaloes.” GigaScience 9 (2): giz166.

Suzuki, Nodoka, Alexei V. Abramov, Yosuke Amaike, Yoshinori Nishita, and Ryuichi Masuda. 2023. “Phylogeography of the Altai Weasel (Carnivora: Mustelidae: Mustela Altaica) Based on an Analysis of Mitochondrial Control-Region Haplotypes.” Biological Journal of the Linnean Society. Linnean Society of London 138 (3): 274–81.

Swali, Pooja, Rick Schulting, Alexandre Gilardet, et al. 2023. “Yersinia Pestis Genomes Reveal Plague in Britain 4000 Years Ago.” Nature Communications 14 (May): 2930.

Torgerson, Dara G., Rob J. Kulathinal, and Rama S. Singh. 2002. “Mammalian Sperm Proteins Are Rapidly Evolving: Evidence of Positive Selection in Functionally Diverse Genes.” Molecular Biology and Evolution 19 (11): 1973–80.

Tzedakis, Polychronis C., David A. Hodell, Christoph Nehrbass-Ahles, Takahito Mitsui, and Eric W. Wolff. 2022. “Marine Isotope Stage 11c: An Unusual Interglacial.” Quaternary Science Reviews 284 (May): 107493.

Valk, Tom van der, Patrícia Pečnerová, David Díez-del-Molino, et al. 2021. “Million-Year-Old DNA Sheds Light on the Genomic History of Mammoths.” Nature 591 (7849): 265–69.

Van Valen, Leigh. 1976. “Ecological Species, Multispecies, and Oaks.” Taxon 25 (2/3): 233–39.

Vasiliev, S. K. 2021. “Remains of the Baikal yak (Poehpagus mutus baikalensis N. Verestchagin, 1954) from Late Pleistocene localities of Southern Siberia.” Trudy Zoologicheskogo instituta 325 (4): 384–408.

Verdugo, Marta Pereira, Victoria E. Mullin, Amelie Scheu, et al. 2019. “Ancient Cattle Genomics, Origins, and Rapid Turnover in the Fertile Crescent.” Science 365 (6449): 173–76.

Verestchagin, N. K. 1954. “Baikal Yak (Poёphagus Baikalensis n. Ver., Sp. Nova, Mammalia) from the Pleistocene Fauna of Eastern Siberia.” Doklady Akademii Nauk SSSR 99 (3): 455–59.

Wang, Kun, Lizhong Wang, Johannes A. Lenstra, et al. 2017. “The Genome Sequence of the Wisent (Bison Bonasus).” GigaScience 6 (4): 1–5.

Wang, Wei, Alim Samat, Yongxiao Ge, et al. 2020. “Quantitative Soil Wind Erosion Potential Mapping for Central Asia Using the Google Earth Engine Platform.” Remote Sensing 12 (20): 3430.

Wang, Xin, Haitao Wei, Mehdi Taheri, Farhad Khormali, Guzel Danukalova, and Fahu Chen. 2016. “Early Pleistocene Climate in Western Arid Central Asia Inferred from Loess-Palaeosol Sequences.” Scientific Reports 6 (1): 20560.

Węcek, Karolina, Stefanie Hartmann, Johanna L. A. Paijmans, et al. 2017. “Complex Admixture Preceded and Followed the Extinction of Wisent in the Wild.” Molecular Biology and Evolution 34 (3): 598–612.

Wei, Caihong, Huihua Wang, Gang Liu, et al. 2016. “Genome-Wide Analysis Reveals Adaptation to High Altitudes in Tibetan Sheep.” Scientific Reports 6 (1): 26770.

Wu, Dong-Dong, Xiang-Dong Ding, Sheng Wang, et al. 2018. “Pervasive Introgression Facilitated Domestication and Adaptation in the Bos Species Complex.” Nature Ecology & Evolution 2 (7): 1139–45.

Wu, Gujie, Hefei Ren, Qin Hu, et al. 2023. “The Circadian Rhythm Key Gene ARNTL2: A Novel Prognostic Biomarker for Immunosuppressive Tumor Microenvironment Identification and Immunotherapy Outcome Prediction in Human Cancers.” Frontiers in Immunology 14 (May): 1115809.

Wu, Xinming, Lu Xu, Haoyuan Zhang, et al. 2024. “Genome-Wide Selection Sweep Analysis to Identify Candidate Genes with Black and Brown Color in Tibetan Sibu Yaks.” Animals: An Open Access Journal From MDPI 14 (17): 2458.

Xu, Jianfeng, Yanyan Wang, Fuyue Shi, et al. 2025. “Genomic Diversity and Selection Signatures for Zaosheng Cattle.” Biology 14 (6): 623.

Xuebin, Q., H. Jianlin, B. Lkhagva, et al. 2005. “Genetic Diversity and Differentiation of Mongolian and Russian Yak Populations.” Zeitschrift Für Tierzuchtung Und Zuchtungsbiologie [Journal of Animal Breeding and Genetics] 122 (2): 117–26.

Yang, Lin, Hong Jin, Qien Yang, et al. 2025. “Genomic Evidence for Low Genetic Diversity but Purging of Strong Deleterious Variants in Snow Leopards.” Genome Biology 26 (1): 94.

Yang, Tianfu, Michelle Miller, David Forgacs, James Derr, and Paul Stothard. 2020. “Development of SNP-Based Genomic Tools for the Canadian Bison Industry: Parentage Verification and Subspecies Composition.” Frontiers in Genetics 11 (November). 10.3389/fgene.2020.585999.

Yu, Jianhua, Wei Cong, Xiuming Li, Lu Wang, Kun Jin, and Yuguang Zhang. 2025. “Whole-Genome Resequencing Reveals Population Genetic Structure and Selection Signatures in the Golden Wild Yak.” Diversity 17 (10): 687.

Yun, Yan, Xiaxin Yang, Shichuan Tan, Pin Wang, Yanbin Ji, and Xiulian Sun. 2022. “Targeting Upregulated RNA Binding Protein RCAN1.1: A Promising Strategy for Neuroprotection in Acute Ischemic Stroke.” CNS Neuroscience & Therapeutics 28 (11): 1814–28.

Zhang, K., J. A. Lenstra, S. Zhang, W. Liu, and J. Liu. 2020. “Evolution and Domestication of the Bovini Species.” Animal Genetics 51 (5): 637–57.

Zhang, Shangzhe, Wenyu Liu, Xinfeng Liu, et al. 2021. “Structural Variants Selected during Yak Domestication Inferred from Long-Read Whole-Genome Sequencing.” Molecular Biology and Evolution 38 (9): 3676–80.

