## Supplementary Information and Figures for "Ancient DNA resolves the taxonomic status and functional characteristics of the Pleistocene lowland yak (*Bos baikalensis*)"

### *Morphological assignment to Baikal yak*

The limb bones of the Baikal yak differ from those of steppe bison (*Bison priscus*), which coexisted in the same region, primarily by Baikal yak having smaller overall dimensions. Most of the Baikal yak limb's large long bones are noticeably shorter and relatively more robust than those of the bison, with the metacarpals being the most diagnostic elements. The metacarpals of the Baikal yak include a more flattened diaphysis and articular condyles at the distal end that are more elongated in the lateromedial direction than in steppe bison. Unlike in steppe bison, the tuberosities situated above the distal epiphysis of the metacarpal are weakly developed or entirely absent in the Baikal yak. The distal humerus clearly distinguishes the Baikal yak from steppe bison on the basis of structural features, such as a shorter articular trochlea and a relatively shorter and higher medial lip of the trochlea in the Baikal yak. For the two morphologically identified Baikal yak specimens examined in this study, the diagnostic measurement for the metacarpal is its overall length (L; 205.2 mm), while the humerus was best distinguished by its length from the head (GLC; 317 mm) and diameter of the upper end (Dp; 125 mm) (Vasiliev 2021) (Table S1).

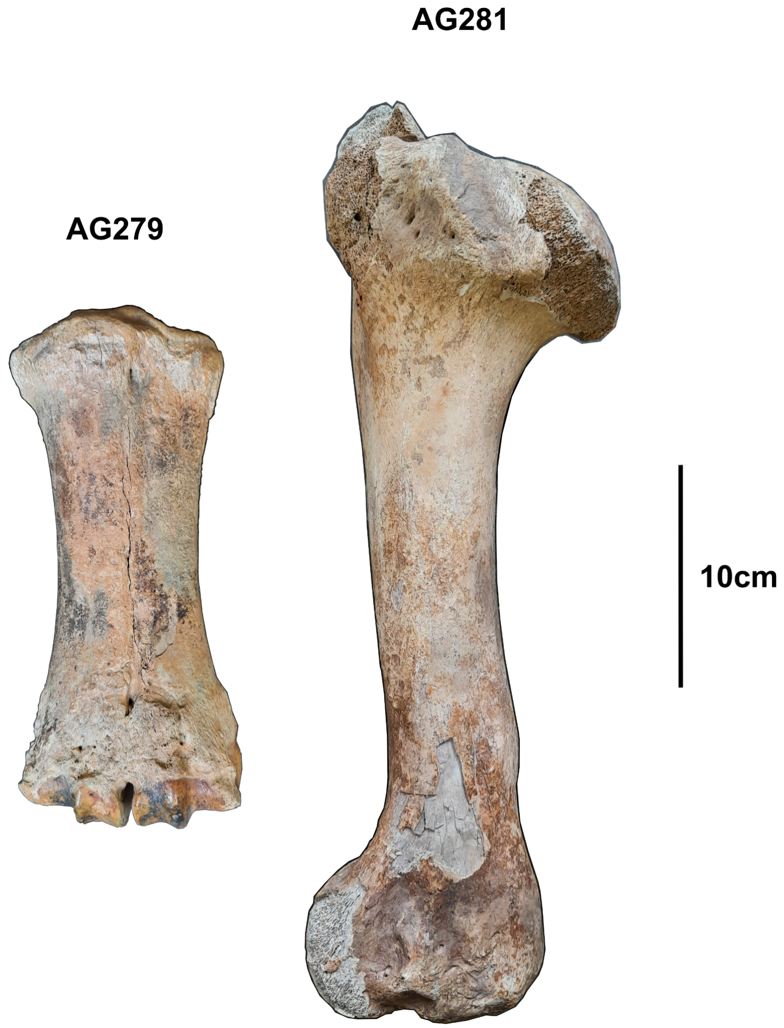


***Figure S1: Other sides for the two Baikal yak voucher specimens from Fig. 1A.*** *Scale is shown.*

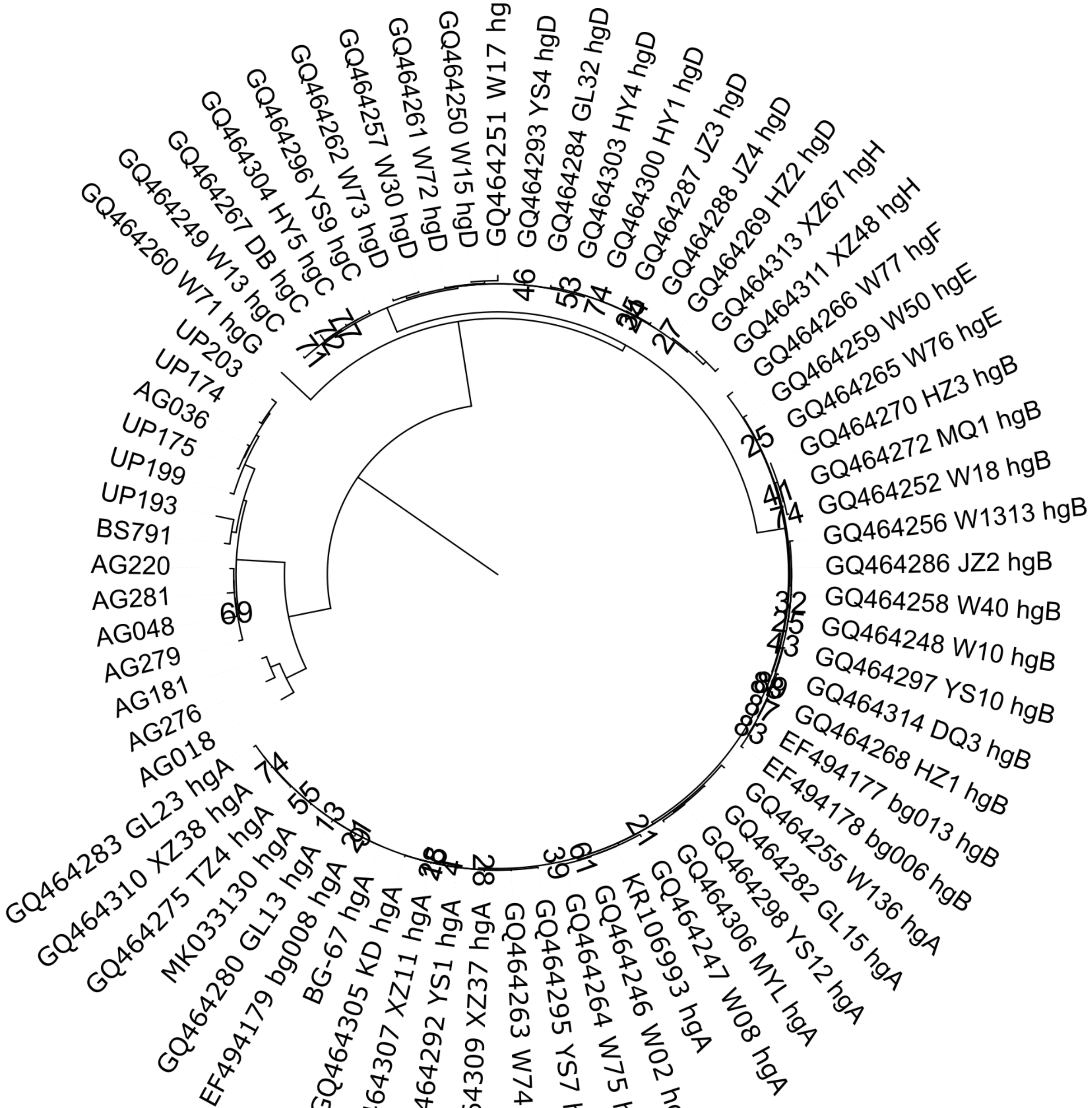


***Figure S2: Mitochondrial phylogeny from Fig. 1B presented with node support values below 90%.*** *Individual IDs are shown, hg stands for haplogroup.*

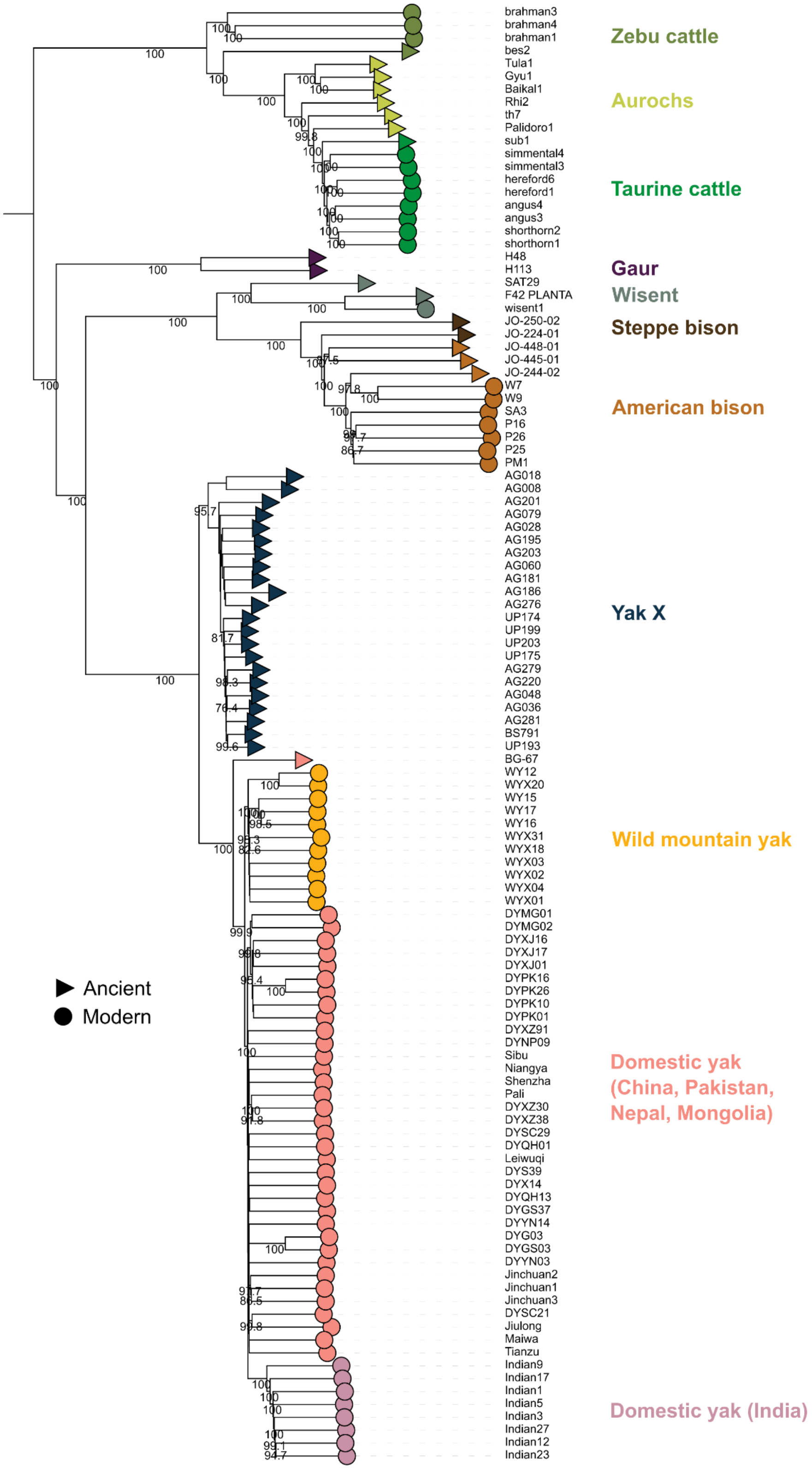


***Figure S3: Autosomal phylogeny from Fig. 1C presented with branch support values above 75%.*** *Individual IDs are shown.*

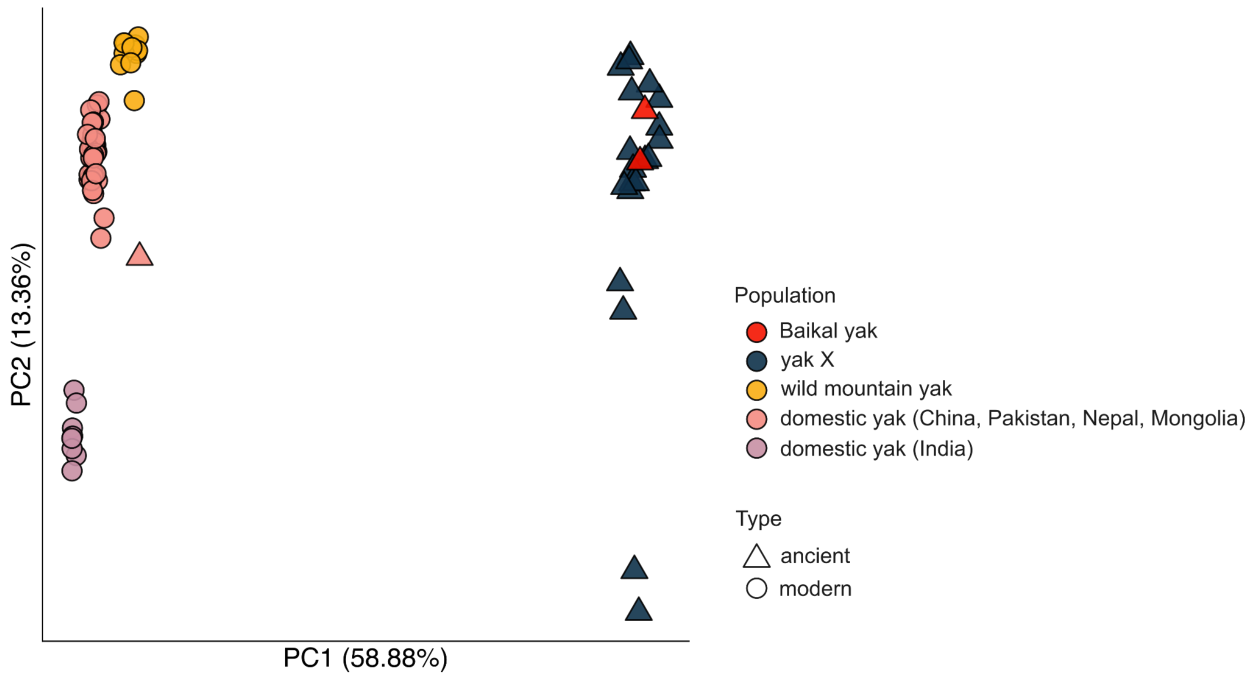


***Figure S4:* f3*-statistics-based PCA containing yak diversity and the two Baikal yak voucher specimens.***

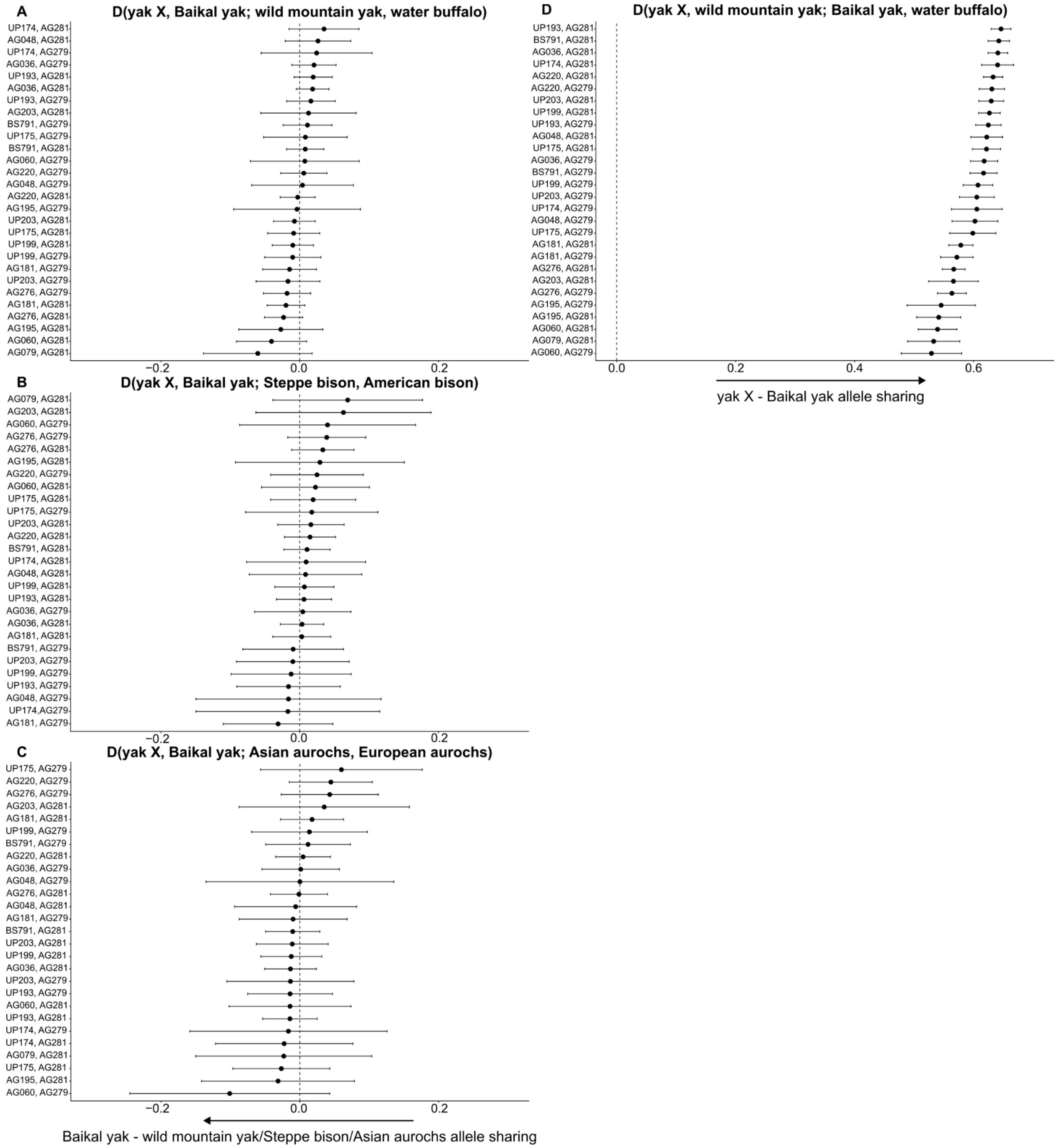


***Figure S5: Allele sharing statistics testing for Baikal yak specimens forming a clade with yak X relative to A) wild mountain yak (WYX20) and water buffalo (gzb1), B) steppe bison (JO-250-02) and American bison (SA3), C) Asian aurochs (Tula1) and European aurochs (Palidoro1).*** *All $-3 \leq Z \leq 3$* ***D) Allele sharing statistics testing for excess affinity between pairs of Baikal yak specimens and yak X relative to wild mountain yak.*** *All $Z > 3$. Error bars show ± 3 standard errors. Pairs of yak X, Baikal yak individuals shown on y-axis.*

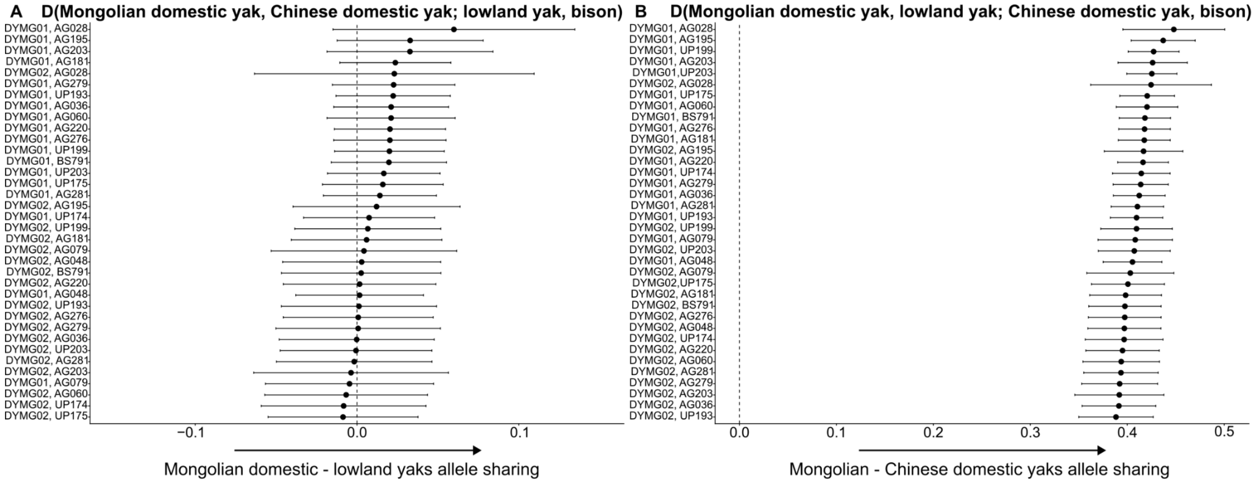


***Figure S6: Allele sharing statistics testing for excess affinity between Mongolian domestic yaks (DYMG01 and DYMG02) and lowland yaks relative to a Chinese domestic yak (Leiwuqi) and bison (SA3).*** *Error bars show ± 3 standard errors. Pairs of Mongolian domestic and lowland yak individuals shown on y-axis. A) All -3 ≤ Z ≤ 3. B) All Z > 3.*

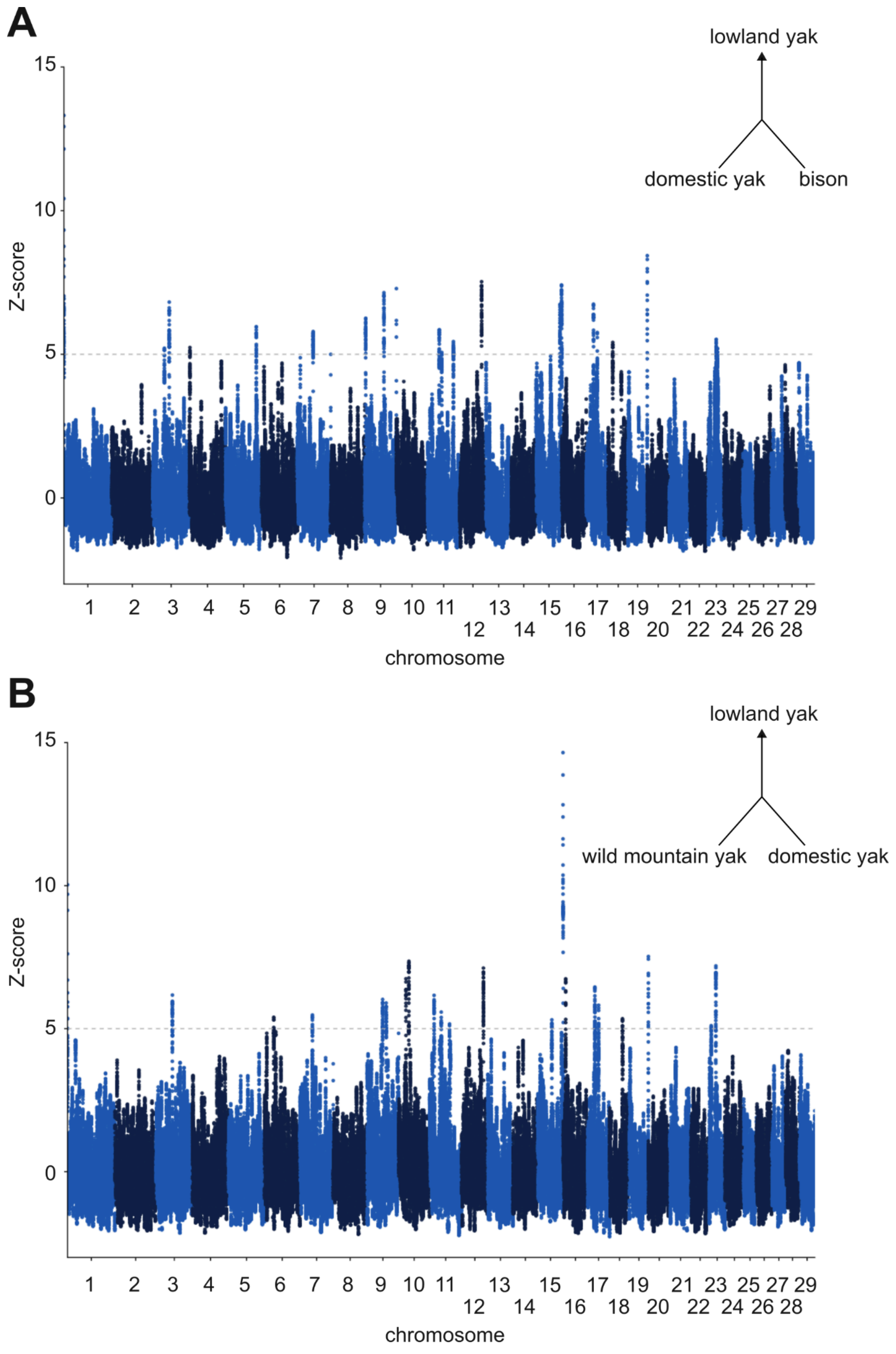


***Figure S7: Alternative versions of the* f3*-statistics-based Population Branch Statistic (PBS) outlier test***, *identifying highly differentiated regions in the lowland yak relative to A) domestic yak and bison or B) wild mountain and domestic yaks, using A) 390 or B) 392 random windows for establishing a null distribution to calculate Z-scores. A schematic of the statistics is depicted.*

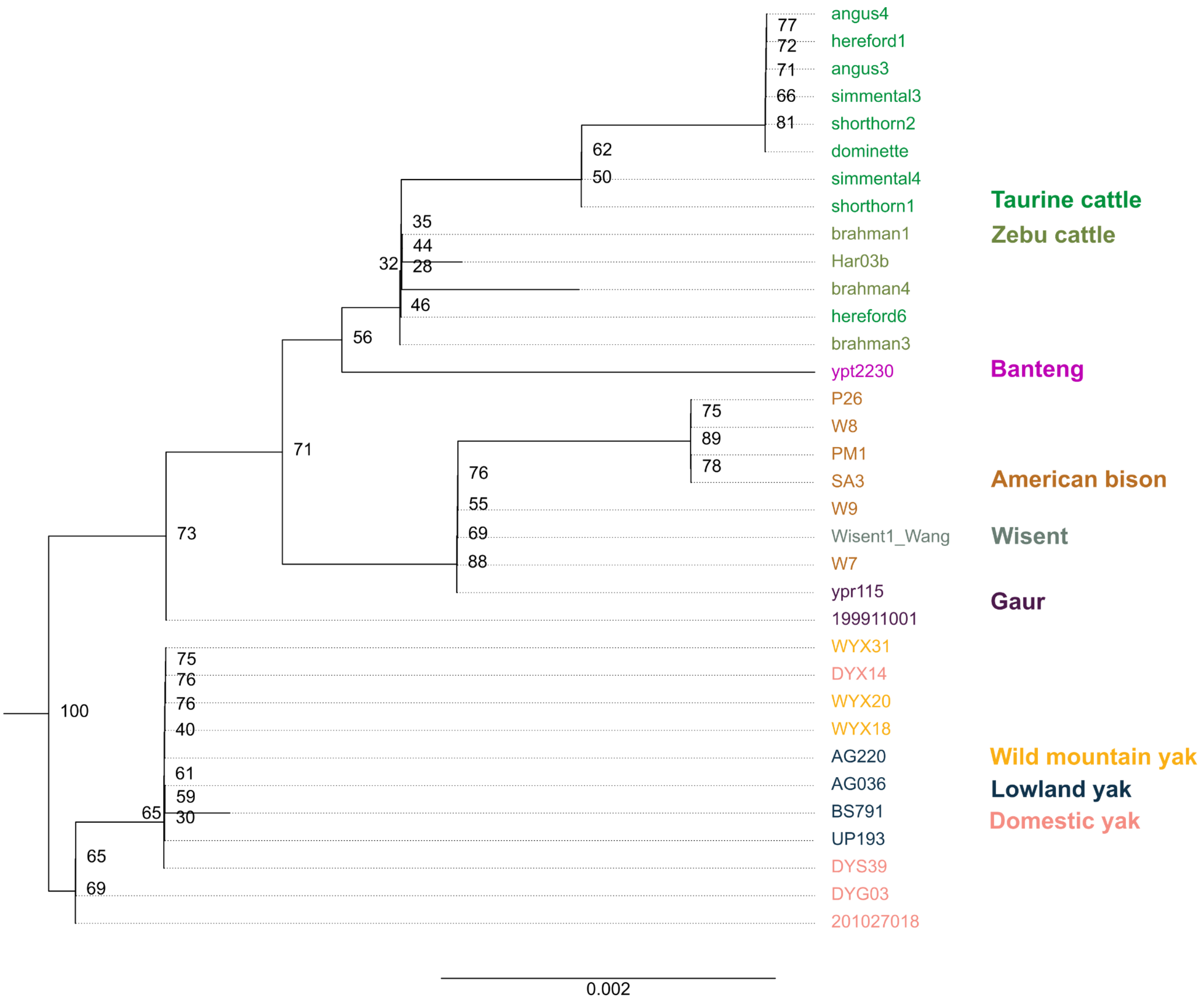


***Figure S8: Maximum-likelihood phylogenetic tree of* EPAS1 *exons across bovine diversity.*** *Node support is shown. Nucleotide substitution per site is shown as scale. The tree was estimated from 2,882 sites, using a HKY+F model, and rooted with water buffalo (not shown).*

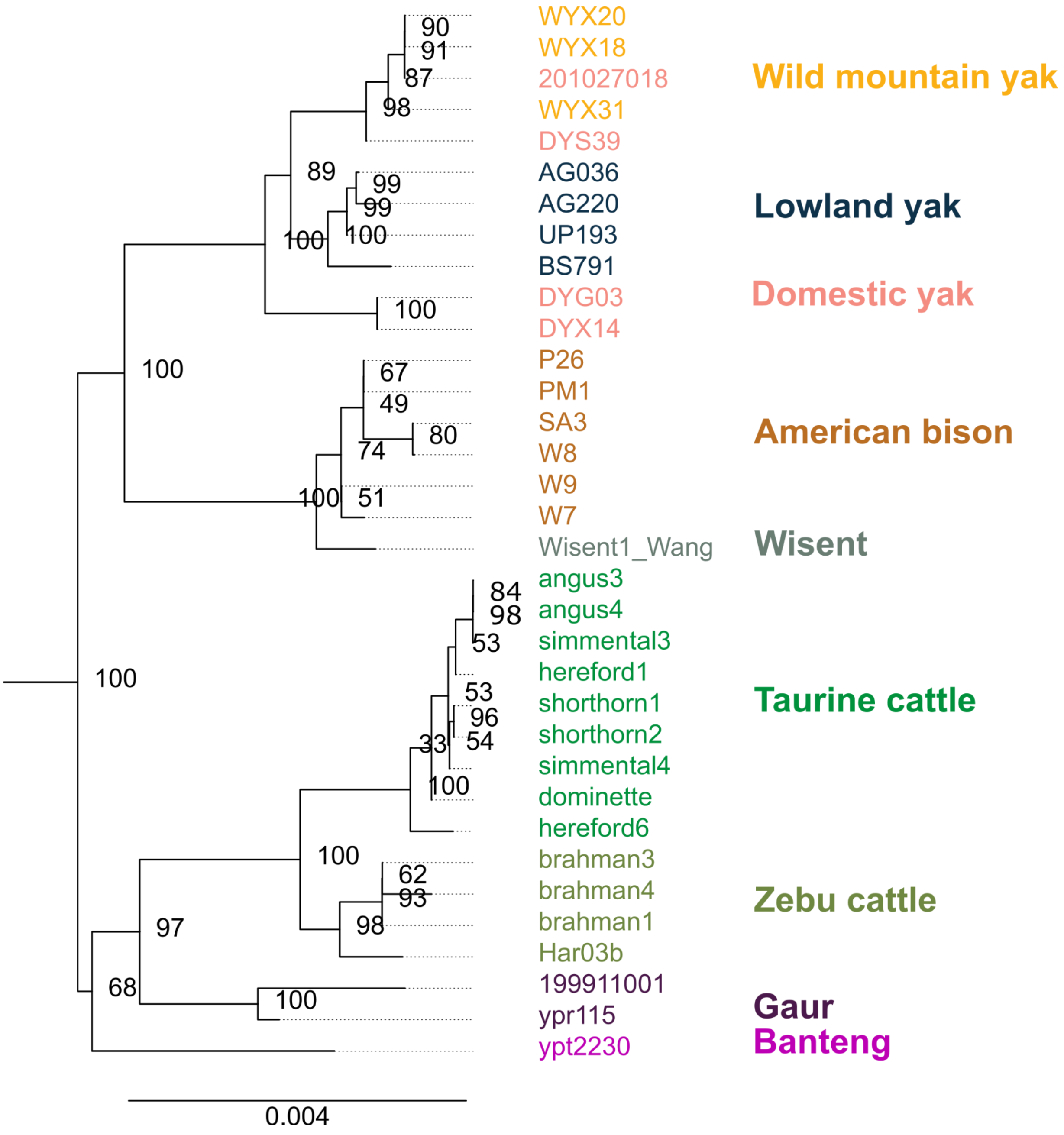


***Figure S9: Maximum-likelihood phylogenetic tree of the whole* EPAS1 *gene (chr11:28,735,330-28,825,665) across bovine diversity.*** *Node support is shown. Nucleotide substitution per site is shown as scale. The tree was estimated from 48,065 sites using a TIM2+F+R3 model, and rooted with water buffalo (not shown).*
